# A Common Reference Landscape of Mouse CD4+ T Cell States

**DOI:** 10.64898/2026.05.29.728553

**Authors:** Nitya Mehrotra, Derek Bangs, Andrea Lebron Figueroa, Gavyn Chern Wei Bee, Rocky Lai, Tomoyo Shinkawa, Kevin C. Osum, Antoine Freuchet, Ziang Zhang, Lilian Nogueira, Taylor Heim, Ken Cadwell, Michael Starnbach, Samuel Behar, Stefani Spranger, Marion Pepper, Marc K. Jenkins, Julia Merkenschlager, Christophe Benoist, David Zemmour, the immgenT Project

## Abstract

CD4+ T cells are orchestrators of the immune system with diverse effector functions. Their full molecular diversity remains unclear due to the lack of a unified framework. Within the immgenT project, we profiled RNA, surface markers, and TCR clonotypes in conventional CD4+ T cells across >700 samples. Integration with a joint RNA-protein deep generative model revealed an ensemble of 20 CD4 states that account for all cells across tissues and challenges. Small, highly polarized clusters that evoke Th1, Th2, Th17, and Tfh states co-exist with a majority of activated cells with mixed programs. Unlike CD8+ T cells, memory CD4+ cells largely mapped to the same states as effectors. Resting states proved quite diverse, and we uncovered unexpected similarities between Tfh and chronically-stimulated states. Together, immgenT provides a unified molecular reference for CD4+ T cell diversity.

## Introduction

CD4+ T cells are arguably the principal regulators of the immune system, coordinating inflammation, enforcing tolerance, and orchestrating humoral immunity. These cells diverge fundamentally from cells of the CD8+ lineage because their T cell antigen receptors (TCRs) recognize peptides bound to Major Histocompatibility Complex class II (MHCII) molecules^1–3^ and provide “helper” functions that augment the antibody- and cytotoxicity-based functions of the adaptive immune system, and orchestrate innate cell differentiation^4–7^. Consequently, CD4+ T cell deficiencies lead to lethal opportunistic infections if left untreated^8,9^, and CD4+ T cells drive a variety of allergic and autoimmune pathologies. Their broader extra-immunological roles are also being recognized^10–12^.

Although some functions of CD4+ T cells are actuated through cell-cell contact, many of their effector functions are mediated by the wide array of cytokines and chemokines they can produce, each of which has major pleiotropic effects on other immune cells: IL-2 regulating growth and differentiation, IFNg promoting innate immune cell function and defenses against intracellular pathogens, IL-4 and IL-21 driving B cell differentiation and immunoglobulin affinity maturation, IL-17 orchestrating defenses against extracellular bacterial and fungal pathogens, IL-5 and IL-13 driving protective responses to helminth infections or inducing allergic diseases, and IL-10 as a broad anti-inflammatory mediator^7^. Accordingly, much of the framework around classification of CD4+ T cells is founded upon these cytokines and the transcription factors that drive their expression, starting with the seminal distinction between Th1 and Th2 T cells^13^, and expanding to Th17 and other specialized subsets^7,14–18^. Indeed, the 1/2/17 organization has been applied to other lineages, such as innate lymphocytes^19^, but not to CD8+ T cells, whose partitioning is organized around memory and residence concepts^20^.

On the other hand, the simple subsetting of CD4+ T cells into discrete “Th” lineages has long been questioned as overly simplistic, as this compartmentalization ignores multi-functional T cells or functional plasticity, leading to sometimes erroneous tagging of diseases as “Thx pathologies”^21–25^. In addition, single-cell transcriptomics has often failed to find *in vivo* the equivalent of the discrete Th states that can be induced by cytokine cocktails *in vitro*, instead describing overlapping functional continua^26–28^. These studies led to a perspective in which states of CD4+ T cells *in vivo* fall along polar continua rather than discrete subsets, with mixed transcriptional phenotypes arising from non-exclusive combinations of cytokine modules.

We revisited the question of CD4+ T cell diversity in the context of the immgenT program^29^. This multi-participant project generated integrated single-cell transcriptomic and surface-protein (128-marker panel) profiling of >750,000 T cells from 734 samples spanning 45 different tissues and conditions (infectious, autoimmune, tumoral). We focus here on *Foxp3*- conventional CD4+ T cells (T regulatory and non-conventional CD4+ T cells are analyzed in a companion manuscript). We resolve 20 robust CD4+ T cell states, “gravitational poles” that capture resting and antigen-experienced states across organs and conditions, and describe a graded polarization model of CD4+ T cell differentiation, in which a minority of cells are pushed into polarized extremes, coexisting with a broader pool of activated but less committed cells, within which Th programs also exist, though less univocally.

The CD4 immgenT framework provides a comprehensive single-cell reference system for examining mouse CD4+ T cell differentiation, with online tools for detailed expression of genes and proteins in the immgenT data, or for projecting external data into it (immgenT reference-based integration, T-RBI), and cytometry marker panels to relate routine flow cytometry to the immgenT framework.

## Results

### Scope and overview of CD4+ T cells in immgenT

As part of the immgenT effort to comprehensively analyze mouse T cell states, 243,153 single CD4⁺ T cells (hereafter “CD4”) were profiled across 734 samples. In addition to transcriptome (scRNAseq), paired αβTCR (TCRseq) and expression of 128 surface protein markers were assessed on the 10X Genomics single-cell platform. These data span over 46 anatomical locations and 127 distinct experimental conditions (**Extended Data Table 1**). In addition to secondary lymphoid organs (SLO), 37% of cells originated from non-lymphoid tissues, and 55% from settings of immunological perturbation (infection, autoimmunity, immunization, cancer). Some datasets included CD4+ T cells of known specificity, identified by transfer of cells from TCR-transgenic mice or by peptide-MHC (pMHC) tetramer staining. This enabled direct comparison of antigen-specific CD4+ T cells across tissues and at different phases of an immune response.

As detailed in the companion immgenT-Cosmology manuscript^29^, all T cells were integrated using totalVI^30^, a deep learning framework that jointly models RNA and protein measurements to define a shared latent space, and a unified dimensionality reduction (the “all-T Minimal Distortion Embedding (MDE)”, **Fig. 1a**). CD4 cells formed distinct compartments, clearly separated from the other seven T cell lineages, including CD4⁺Foxp3⁺ regulatory T cells. To better visualize fine heterogeneity within CD4+ T cells, we computed a CD4-specific MDE and defined 20 clusters that were robust across both RNA and protein modalities (robustness and generalizability are detailed in the accompanying immgenT-Cosmology^29^) (**Fig. 1b, Extended Data Table 2**). This framework captures the diversity of CD4+ T cell states within each sample while aligning shared states across conditions. Within it, the main distinction was between groups of “resting” (A-F) and “activated” (G-Y) clusters, the cell-surface CITEseq values matching the classic CD62L/CD44-based distinction (**Fig. 1c**) (a small number of CD44⁺ cells were present in the resting clusters, which included some memory cells (**Fig. S1a**)). Cluster P represents cycling cells, and three additional “workshop status” clusters, CD4.wM, CD4.wN, and CD4.wZ, whose reliability is uncertain, are discussed in the Legend.

**Figure 1.**
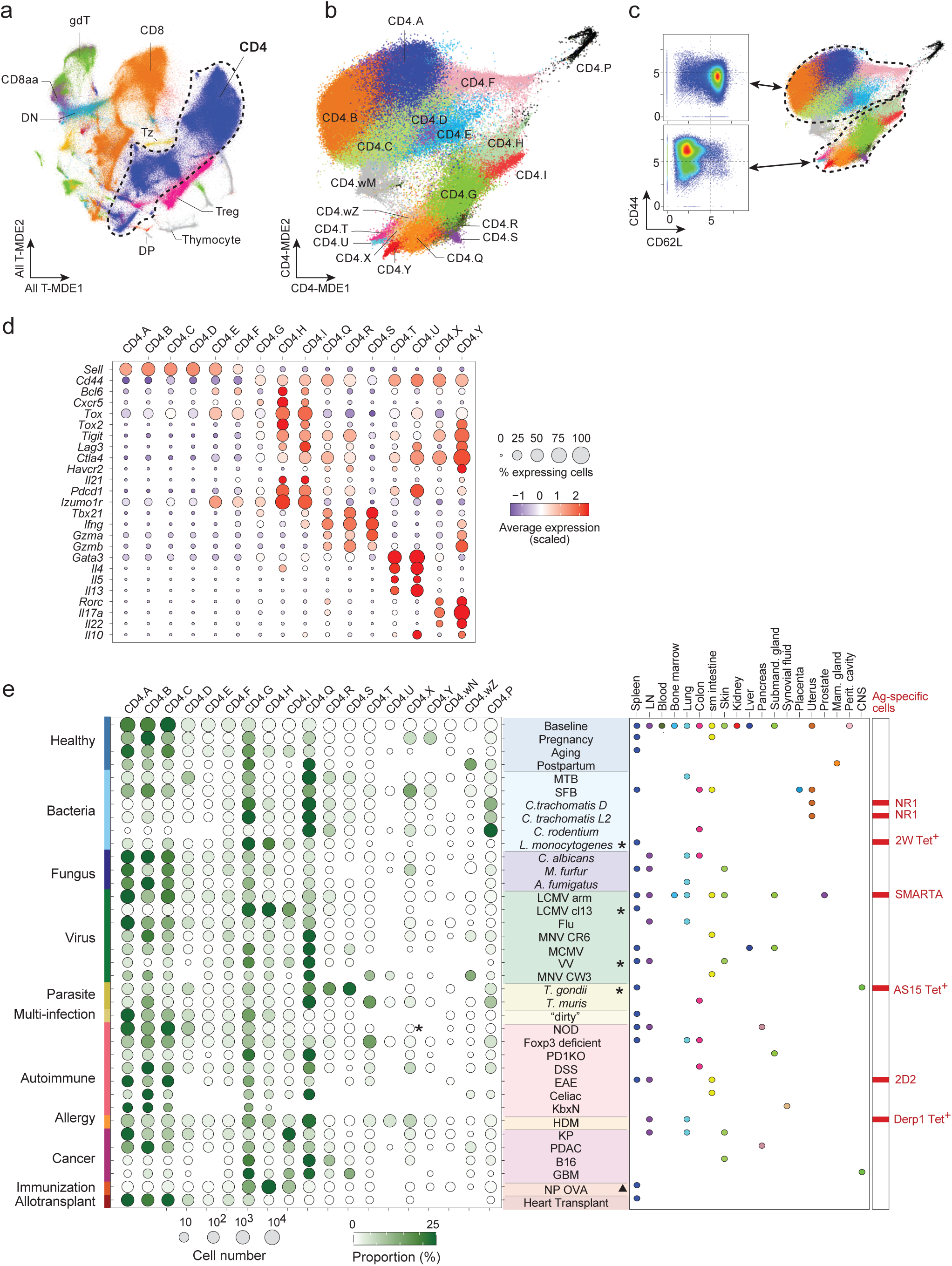
immgenT resolves CD4+ T cell heterogeneity into 20 discrete clusters and a reference embedding. **(a)** Minimum Distortion Embedding (MDE) of all T cells (all-T MDE) profiled in the immgenT dataset, colored by major T cell lineages: CD4, CD8, CD8αα, DN, DP, γδ T cells, Zbtb16⁺ T cells (Tz; including iNKT and MAIT cells), Treg, and thymocytes. CD4 (CD8⁻ CD4⁺ Foxp3⁻ conventional αβ T cells) comprised 243,153 of 682,951 total T cells profiled across tissues and immune conditions using 10x Genomics 5′ v2 single-cell RNA sequencing with paired CITE-seq (128-plex) and αβ TCR sequencing. **(b)** CD4-specific MDE colored by cluster identity. Cluster labels prefixed with “w” denote provisional clusters, typically small (<1% for wN and wZ), and are not further analyzed in this study. **(c)** Flow cytometry–like representation of CD62L versus CD44 expression (CITE-seq; log1p(CP10K)) across resting (A–F) and activated (G–Y) clusters. **(d)** Dot plot showing expression of canonical CD4 differentiation genes across clusters. Color indicates average expression (Z-score), and dot size represents the fraction of cells expressing each gene. **(e)** Overview of the immgenT dataset (detailed descriptions are provided in **Extended Data Table 1**). For each immune condition (rows), the left dot plot shows the number of cells (dot size) and the mean cluster proportion (color). When a specific population was profiled, a symbol is shown next to the legend: a triangle denotes CXCR5⁺ PD1⁺ cells and a star denotes CD44⁺ cells. The right panels indicate the organs profiled and whether antigen-specific cells were included. NR1, *Chlamydia*-specific transgenic cells; SMARTA, LCMV-specific transgenic cells; 2D2, MOG-specific transgenic cells; 2W tet⁺, antigen-specific cells responding to *Listeria monocytogenes* engineered to express the 2W peptide; AS15 tet⁺, antigen-specific cells responding to *Toxoplasma gondii*. Abbreviations: MTB: Mycobacterium tuberculosis; SFB: segmented filamentous bacterium; C. trachomatis: Chlamydia trachomatis; C. rodentium: Citrobacter rodentium; L. monocytogenes: Listeria monocytogenes; C. albicans: Candida albicans; M. furfur: Malassezia furfur; A. fumigatus: Aspergillus fumigatus; LCMV arm: lymphocytic choriomeningitis virus Armstrong; LCMV cl13: lymphocytic choriomeningitis virus clone 13; Flu: influenza virus; MNV: murine norovirus; MCMV: murine cytomegalovirus; VV: vaccinia virus; T. gondii: Toxoplasma gondii; T. muris: Trichuris muris; “dirty”: dirty-mouse or diverse microbial exposure condition; NOD: non-obese diabetic; PD1KO: programmed cell death protein 1 knockout; DSS: dextran sulfate sodium; EAE: experimental autoimmune encephalomyelitis; Celiac: celiac disease model; KbxN: K/BxN arthritis model; HDM: house dust mite; KP: Kras G12D-driven, p53-deficient lung tumor model; PDAC: pancreatic ductal adenocarcinoma; B16: B16 melanoma model; GBM: glioblastoma; NP-OVA: nitrophenyl–ovalbumin immunization; Heart Transplant: cardiac allotransplantation model.

Genes commonly associated with CD4+ T cell differentiation and function were sharply demarcated in their distribution across clusters (**Fig. 1d**). The expression of chemokine receptors varies widely, to the point where some have used them as proxies of T cell states^31,32^. However, the strikingly variegated patterns of chemokine receptor expression (**Extended Data Fig. 1b**) indicate that no single chemokine receptor can serve as a proxy for any particular cluster.

No cluster derived from one particular immune challenge or tissue (**Fig. 1e)**. Resting clusters were enriched in SLOs and steady-state conditions, while the continuum of activated or antigen-experienced CD44⁺CD62L⁻ cells were enriched in perturbed settings. Some of these activated CD4 clusters were enriched in some conditions, like clusters T and U enriched in house-dust mite (HDM) and *T. muris* models, or clusters X and Y associated with SFB colonization and fungal infection (*M. furfur*).

Antigen-specific cells mapped to the same transcriptional landscape (**Fig. 2a**). In datasets lacking antigen-specific tracking, endogenous responses could instead be identified through expanded paired αβ TCR clonotypes^33–35^ (large expansions with at least 10 cells per clonotype; **Fig. 2b, Extended Data Fig. 2a, Extended Data Table 3**). These endogenous clonal responses closely mirrored tracked antigen-specific populations; for example, the dominant clonotype during acute LCMV Armstrong closely resembled SMARTA cells and expanded clonotypes during *C. trachomatis* infection mapped similarly to NR1 transgenic cells. In several settings, including HDM, SFB, and *M. tuberculosis*, distinct expanded clonotypes occupied non-overlapping regions of the landscape, consistent with reports^36–38^ that TCR specificity influences molecular states, but also suggesting that individual TCR-transgenic models may not capture the full range of antigen-specific responses.

**Figure 2.**
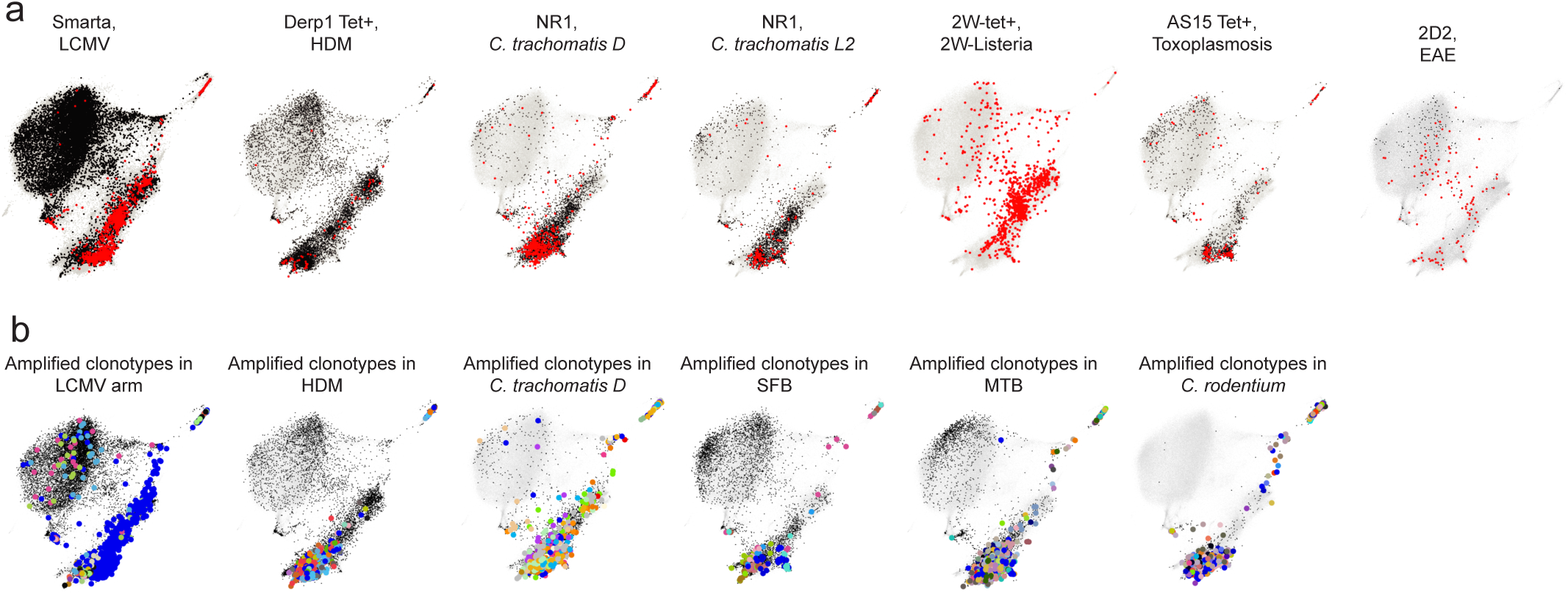
Mapping antigen-specific and clonally expanded cells across immune challenges. **(a)** CD4-MDEs showing the distribution of antigen-specific CD4+ T cells in LCMV Armstrong (SMARTA), HDM (Der p 1 tetramer⁺), C. trachomatis D and L2 (NR1), 2W-L. monocytogenes (2W tetramer⁺), Toxoplasma gondii (AS15 tetramer⁺), and EAE (2D2). Antigen-specific cells are highlighted in red; all other CD4+ T cells from the corresponding condition are shown in black. Background immgenT CD4 cells are shown in grey. **(b)** CD4-MDE showing clonally expanded CD4⁺ T cells (clonotypes represented by more than 10 cells) in LCMV Armstrong, HDM, SFB, C. trachomatis D, MTB, and C. rodentium. Transgenic cells were excluded from the analysis. Cells are colored by clonotype; the remaining CD4+ T cells from each condition are shown in black. Background immgenT CD4 cells are shown in grey.

Beyond serving as a collection of single-cell datasets, the immgenT framework enables mapping of external datasets through T cell Reference-Based Integration (T-RBI), a reference-mapping approach based on scVI and scANVI^39,40^ described and validated in the companion immgenT-Cosmology manuscript^29^. This approach enabled projection of antigen-specific T cells from immune settings not represented in the current dataset, including GP66-specific CD4⁺ T cells during chronic LCMV clone 13 infection and in GP66-expressing MC38 tumors^41^, both of which mapped predominantly to the anergic-like cluster CD4.I, as well as alloantigen-specific T cells^42^ from heart transplantation, which occupied distinct locations in graft and draining lymph node samples (**Extended Data Fig. 2b**). More broadly, T-RBI provides a straightforward way to compare cell populations studied by independent investigators. We illustrate this use case with a recent example from one of the authors’ laboratories. A population of IFNγ⁺ CD4⁺ T cells was identified in the colon of four-week-old mice, but it was unclear whether these cells were analogous to a population of T-bet⁺ cells in the small intestine of adult mice annotated as Tr1 cells because of their propensity to produce IL-10 following anti-CD3 stimulation^43^. When the two datasets were mapped onto the CD4 framework using T-RBI, it became immediately clear that they were nearly identical, both consisting predominantly of CD4.Q cells with smaller populations of more differentiated states (**Extended Data Fig. 2c-f**).

### Activated CD4⁺ T cell states are organized along a graded polarization landscape

The group of activated/experienced clusters formed a suggestive disposition in the MDE, wherein a minority of cells in tips adopt sharp and exclusive phenotypes, derived from a larger pool of cells which express less exclusive mixes of programs (**Fig. 3a**). For the key cytokines that define the broad axes of differentiation, cells with high expression were concentrated in small, sharply defined clusters (hereafter “polarized”) that projected from two large central clusters, G and Q (**Fig. 3a,b**), with lower, mixed expression. The polarized clusters matched classical CD4+ T helper expectations: Th2 (T and U), marked by *Il5* and *Il13*; Th17 (X and Y), marked by *Il17* and *Rorc;* and Th1 (R and S), marked by *Ifng* and *Tbx21*. On the opposite side, *Il21* expression defined a Tfh pole (cluster H, also I). Cytokine expression was mutually exclusive in the polarized clusters. Expression of the corresponding transcription factors (TFs) matched these distributions (**Fig. 3c**), with broader distributions for *Gata3* and *Tbx21* (*Gata3* has a broader role in T cells than merely controlling type-2 cytokines). The poles included cells from a number of datasets (**Fig. 1e**), but predominated in settings where the corresponding differentiation program were expected: *T muris* infection for Th2 poles, Segmented Filamented Bacterium (SFB) for Th17 poles, antigen/adjuvant immunization for Tfh poles, and *Toxoplasma* infection for Th1 poles (**Fig. 3d**).

**Figure 3.**
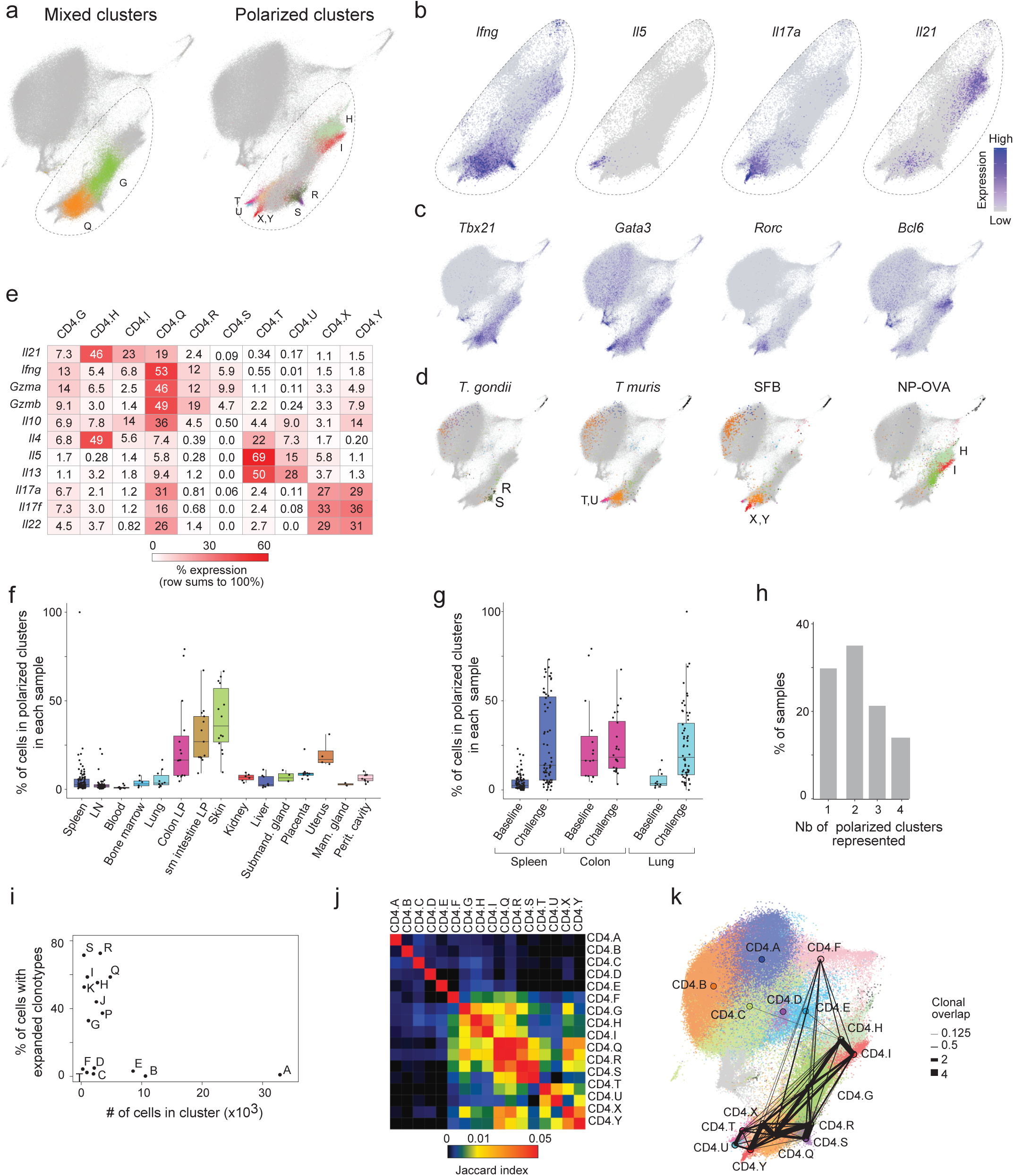
Activated CD4+ T cells map to large mixed T helper phenotype clusters and smaller polarized clusters. **(a)** CD4 MDE highlighting large “mixed-phenotype” clusters (CD4.G, CD4.Q; left) and smaller polarized clusters (right): Th1 (R,S), Th2 (T,U), Th17 (X,Y), and Tfh-like (H,I). Cells are colored by cluster identity; background immgenT CD4+ T cells are shown in grey. **(b)** CD4 MDE showing expression of canonical T helper cytokines (*Ifng, Il5, Il17a, Il21*), zoomed to the activated region shown in (a). **(c)** CD4 MDE showing expression of canonical T helper transcription factors (*Tbx21, Gata3, Rorc, Bcl6*). **(d)** CD4 MDE highlighting representative experimental models: Th1 (*T. gondii* in SLO), Th2 (*T. muris* in colon), Th17 (SFB in ileum), and Tfh (NP–OVA immunization). Cells are colored by cluster identity; background immgenT CD4+ T cells are shown in grey. **(e)** Heatmap showing the contribution of each cluster to total cytokine expression (rows sum to 100%). **(f,g)** Boxplots showing the proportion of cells in polarized clusters (CD4.R, S, T, U, X, Y) among activated CD4+ T cells across organs at baseline (f) and comparing baseline versus immune challenge in spleen, colon, and lung (g). Each point represents one sample. **(h)** Histogram of the number of poles detected per sample (defined as ≥1% of cells). **(i)** Scatterplot of the proportion of cells with expanded clonotypes versus cluster size. **(j)** Heatmap of clonotype sharing between CD4+ T cell clusters (Jaccard index). **(k)** Graph representation of clonotype sharing overlaid on the CD4 MDE. Nodes are positioned at cluster centroids; edge width is proportional to the extent of clonotype sharing. Transgenic cells were excluded from the analysis.

Functionally, cells mapping to polarized clusters disproportionately contributed to cytokine output (**Fig. 3e**). For example, >80% of *Il5* or *Il13* expression localized to Th2 poles (T/U), 70% of *Il17f* to Th17 poles (X/Y). *Ifng* expression was more broadly distributed, with the majority in the mixed clusters G and Q (65%). Together, CD4.G and CD4.Q encompassed the majority of activated/experienced cells (55% of cells; **Fig. 3a**), and showed mixed expression of cytokines and TFs (**Fig. 3e**): some cells in cluster G or Q expressed *Il17a/f*, *Il5*, and the majority of cells in Q expressed *Ifng*. This graded polarization architecture was conserved across tissues and conditions. Polarized clusters were enriched in non-lymphoid tissues (up to 75% in skin at baseline) (**Fig. 3f**) and increased under perturbation (**Fig. 3g**). Most samples contained cells from multiple poles (**Fig. 3h**, **Extended Data Fig. 3a,b**). At baseline, tissues were dominated by G and Q but differed in their representation of individual poles; for example, the gut contained prominent Th17 (X/Y), Th2 (T/U), and Tfh (H) poles; the skin was enriched for Th17 and Th2 poles; and the uterus contained H, S, and T populations (**Extended Data Fig. 3c**). In contrast, spleen and lymph nodes contained relatively few polarized cells (**Extended Data Fig. 3d**), with the notable exception of CD4.H. Consistent with previous reports of steady-state IFNγ-expressing memory-like cells^44^ in SLOs, *Ifng* expression at baseline extended broadly across G–H–I and Q–R–S (**Extended Data Fig. 3e**).

TCR clonotype analysis helped to identify relationships between clusters. First, a high proportion (20-60%) of cells mapping to activated/experienced clusters expressed clonally expanded clonotypes (full nucleotide sequence identity, **Fig. 3i**), whereas expanded clonotypes were predictably very rare in resting clusters. Mapping of overlap between clusters indicated that cells sharing the same TCR clonotype could occupy diverse transcriptional states (**Fig. 3j,k**), although overlap was strongest within polarization groups (H/I, R/S, T/U, X/Y) and between mixed clusters and their closest associated polarized clusters (G with H/I; Q with R/S/T/U/X/Y). Clonal sharing may reflect either a common origin (cells expanding and diverging from the same precursor) or plasticity (cells shuttling between states). In any case, these results imply that CD4+ T responses to a given antigen ultimately generate cells with a variety of functional phenotypes.

Together, these analyses show that qualitative assignment to Th programs is insufficient to describe the full landscape and support a graded model of polarization. In this model, highly polarized, cytokine-producing cells occupy discrete poles, whereas the majority of activated CD4⁺ T cells reside in the central G and Q clusters, apparently expressing lower-intensity, combinatorial helper programs.Helper T cell programs in polarized and mixed clusters: intensity and specificity

These results provided a framework for positioning the classical axes of CD4+ T cell differentiation. The diversity of datasets contributing cells in each state provided a unique opportunity to define the core gene expression programs that characterize these polarized clusters, without the usual biases inherent in one-to-one signature definitions. In other words, by incorporating Type-2 responses from many sources, what is a core Th2 program in vivo? In the less polarized (Q and G): does the mix of programs found there reflect program co-activation in multi-potential cells, or do the cells partition programmatically?

We defined pole-specific gene signatures by comparing each polarized cluster to all others (**Fig. 4a-d, Extended Data Table 4**). Comfortingly, these included classic defining transcripts: *Rorc, Il17a* and *Il23r* for Th17 poles (X and Y)^45–47^, *Gata3, Il4, Il5, Il13* for the Th2 poles^48^, *Cxcr5, Il21* for Tfh poles (H and I)^49,50^ and *Tbx21, Il12rb2* and *Ccl5* for Th1 poles (R and S)^51^. *Ifng* did not meet signature criteria due to its broad expression across multiple clusters. The results uncovered a number of transcripts not commonly associated with those programs, such as *Nmur1*^10,52^ in Th2, members of the Slamf co-receptors (*Slamf2/Cd48* and *Slamf7*)^53^, and S1P receptors (*S1pr5* and *S1pr4*) in Th1, metabolic and structural genes (*Ckb*, *Actn2, Lmnb1*) in Th17. While Th17 pole signature genes are supported by ChIP–seq evidence for canonical regulators, including RORγt, IRF4, BATF, and STAT3^54^. There was little variation in the expression of those genes across tissues at baseline (with some notable exceptions in the expression of canonical cytokines such as *Il4*, *Il21*, and *Il22*) (**Extended Data Fig. 4a-d**).

**Figure 4.**
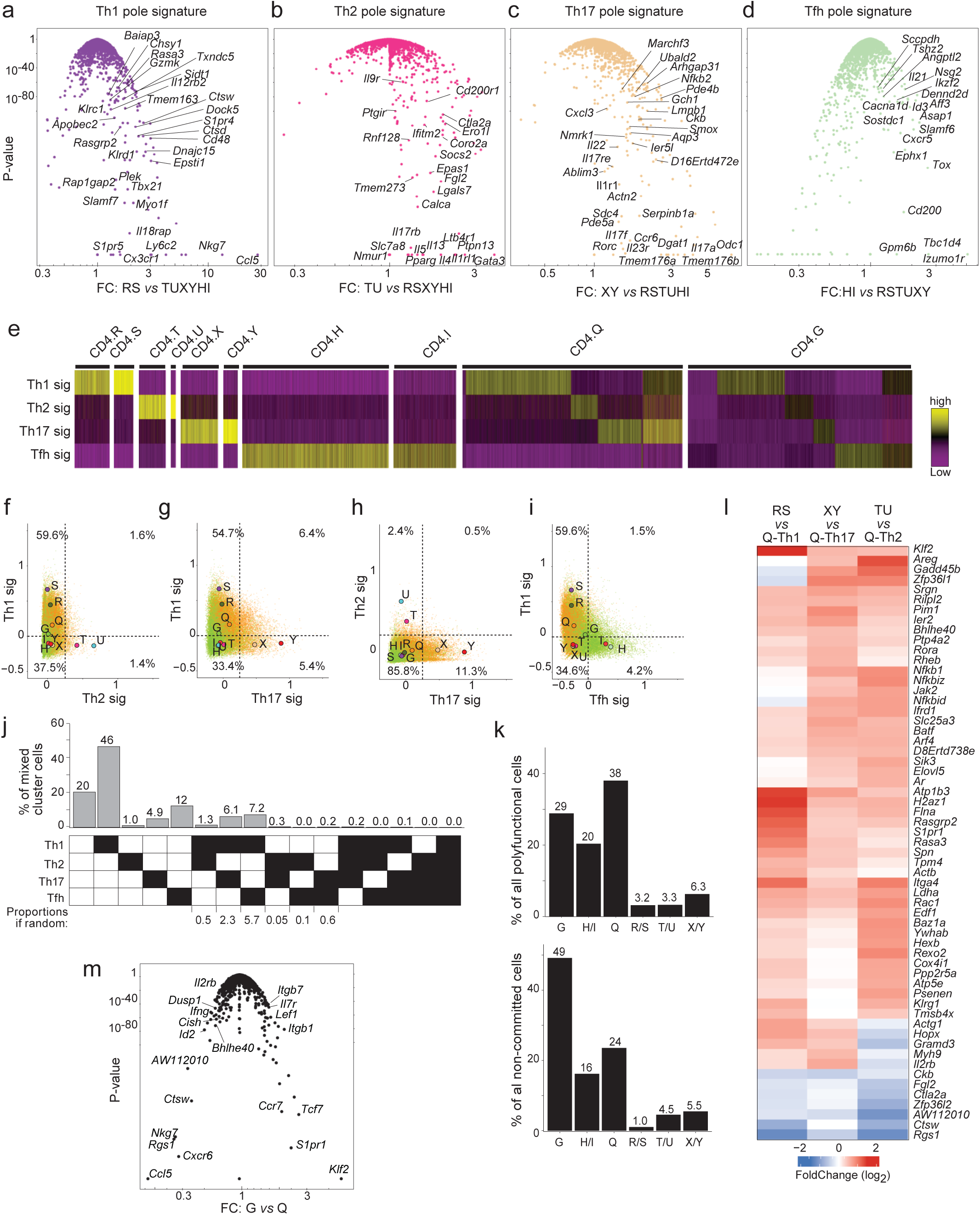
T helper programs vary in intensity and combination across mixed and polarized clusters. **(a–d)** Volcano plots showing differential gene expression between Th1 (CD4.R,S) **(a),** Th2 (CD4.T,U) **(b)**, Th17 (CD4.X,Y) **(c)**, and Tfh (CD4.H,I) **(d)** polarized clusters versus all other poles. Genes are plotted as fold change versus adjusted *P* value. Labeled genes define in vivo Th signatures, selected based on minimum expression (log1p(CP10K) ≥ 0.25), ≥1.5-fold enrichment over the highest-expressing alternative polarized cluster, and adjusted *P* < 10⁻⁵. **(e)** Single-cell heatmap showing expression of Th1, Th2, Th17, and Tfh programs defined in (a–d). Cells downsampled to show at most 10,000 cells per cluster for visualization purposes and ordered by cluster and program expression. **(f–i)** Scatterplots showing pairwise expression of Th programs in CD4.G (green) and CD4.Q (orange) cells. Larger labeled circles indicate the mean program expression of each activated cluster (G–Y). Percentages denote the fraction of CD4.G and CD4.Q cells in each quadrant. **(j)** Distribution of Th program usage in CD4.G and CD4.Q cells. Bars show the percentage of cells with no program, single-program expression, or ≥2 programs (polyfunctional), based on thresholds defined in (f–i). The heatmap below indicates program combinations (black = present), and values above bars denote cell frequencies. Expected frequencies under independence were calculated by multiplying individual program probabilities and are shown beneath the heatmap. **(k)** Distribution of polyfunctional (left) and non-committed (right) cells across CD4 clusters, using thresholds defined in (f–i). **(l)** Heatmap of genes up- and downregulated in each Th polarized cluster relative to the corresponding Th-biased Q population (e.g., RS Th1 pole vs Q-Th1), filtered at FDR < 1% and log2FC > 0.5 in at least two comparisons. **(m)** Volcano plot showing differential gene expression between CD4.G and CD4.Q, controlling for tissue of origin.

To assess the activation of the Th programs at single-cell resolution, we computed integrated scores for each program in every cell. The programs were most prominent in the polarized clusters, where they were mutually exclusive (**Fig. 4e-i, Extended Data Fig. 4e-h**). However, they were also detected in cells from clusters Q and G (**Fig. 4e**). These clusters contained heterogeneous cell populations, including cells with evidence of commitment. Q-Th1, Q-Th2, and Q-Th17 cells retained expression of the corresponding core signatures, albeit at lower levels than cells in the polarized clusters (**Extended Data Fig. 4i-n**). Clusters Q and G also contained polyfunctional cells (**Fig. 4f-k**), most of which expressed both Th1 and Th17 programs in Q (**Fig. 4g**), in agreement with previously described Il17/IFNγ double-expressing cells^26^, or Th1 and Tfh reminiscent of “Tfh1” cells in G^5,55^. **Fig. 4k** quantitatively confirmed that Q and G contained the majority of polyfunctional cells, as well as cells without any of these programs. Thus, the mixed clusters G and Q contain heterogeneous cells with varying levels of “engagement” in final functional programs. These observations prompted us to quantify Th programs and test whether polyfunctional cell frequencies could be explained by the random assortment of individual program activities. In the mixed clusters, polyfunctional frequencies were well predicted by the joint distribution of individual program frequencies (**Fig. 4j**, bottom). In **Extended Data Fig.5**, we formalized this analysis by using the observed marginal probability distribution for each Th program across all cells and computing the expected joint distribution under an independence (random-assortment) model. Comparing expected and observed joint expression as a ratio heatmap showed that, at low Th expression levels corresponding to mixed clusters, the ratio was close to one, indicating good agreement with the model of random combinations. In contrast, polyfunctional cells in the polarized clusters were strongly underrepresented, with ratios approaching zero, corresponding to observed frequencies more than tenfold lower than predicted. These observations imply that cells in the large mixed clusters have not reached final commitment, no stable negative feedback is in place and may activate several of the terminal Th programs at random (with the caveat that we cannot estimate “plasticity”, i.e., how much these patterns fluctuate in a given cell over time). This potential would change in cells from the polarized clusters, where expression of only one program becomes the rule, with negative feedback on the other potentialities. Thus, Th programs inhibit one another as they intensify, resulting in polarized states and more mixed mixed clusters.

**Figure 5.**
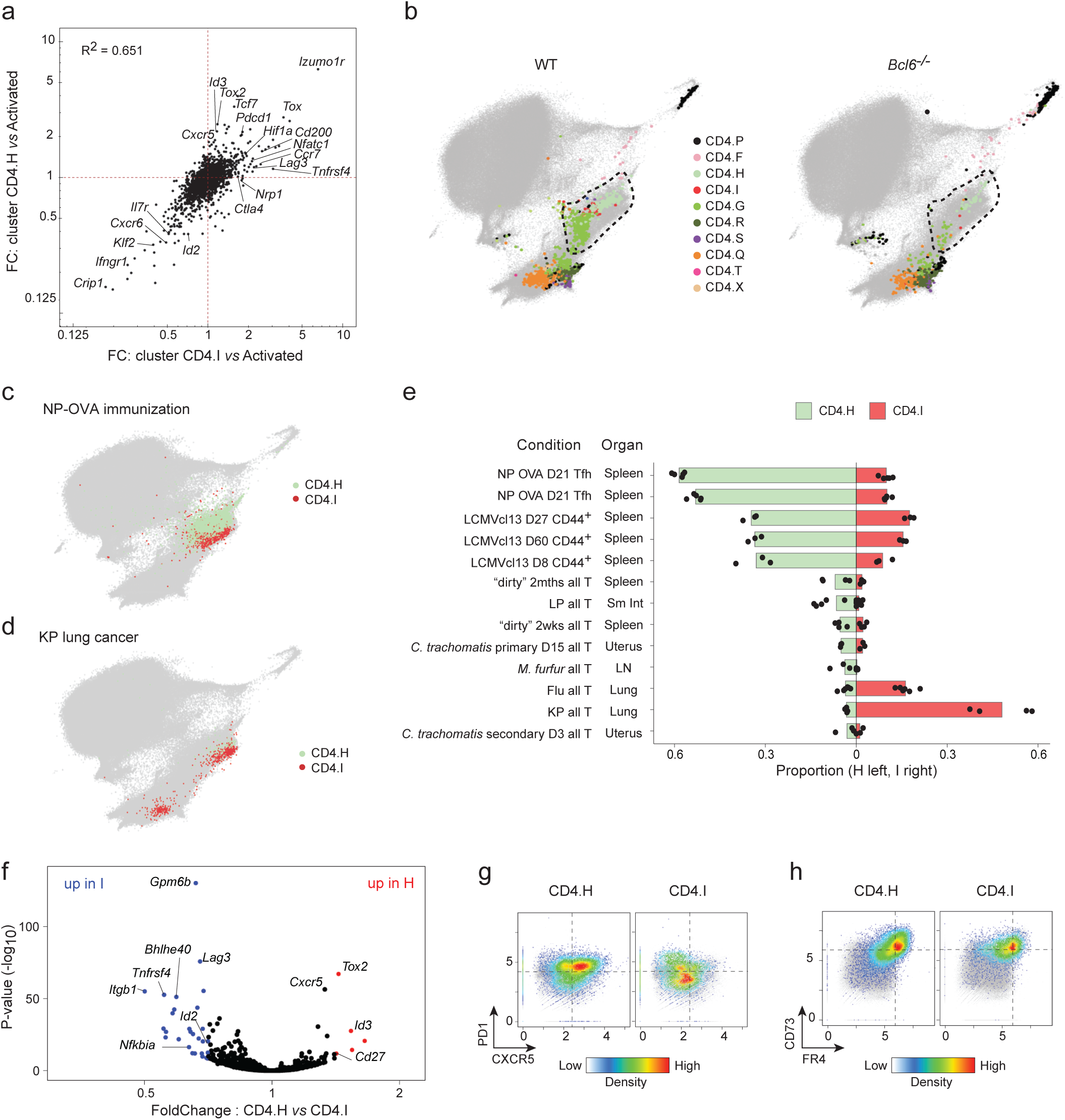
Convergence of Tfh and chronically stimulated CD4+ T cells into CD4.H and CD4.I. **(a)** Fold-change versus fold-change plot comparing gene expression changes in CD4.H and CD4.I (each relative to activated CD4+ T cells). Shared genes lie along the diagonal, whereas genes preferentially enriched in CD4.H or CD4.I fall along the horizontal or vertical axes, respectively. Selected genes are labeled. **(b)** CD4 MDE showing projection of wild-type (left) and Bcl6-knockout (right) LCMV-specific CD4⁺ T cells (SMARTA) at 7 dpi from Nguyen et al.^69^ onto the immgenT CD4 reference using T-RBI (T cell Reference-Based Integration). Cells are colored by cluster identity; background immgenT CD4+ T cells are shown in grey. n = 1870 cells per plot. **(c)** CD4 MDE highlighting CD4.H (light green) and CD4.I (red) cells following NP–OVA immunization; background immgenT CD4+ T cells are shown in grey. **(d)** CD4 MDE highlighting CD4.H (light green) and CD4.I (red) cells infiltrating KP lung tumors; background CD4+ T cells are shown in grey. **(e)** Bar plot showing conditions with the highest proportions of CD4.H (green, left) or CD4.I (red, right). Each dot represents a replicate; bars indicate the mean. **(f)** Volcano plot of differential gene expression between CD4.H and CD4.I, shown as fold change versus adjusted *P* value. Selected genes are highlighted. **(g)** CXCR5 versus PD1 expression in CD4.H and CD4.I (CITE-seq; log1p(CP10K)); color indicates cell density. **(h)** FR4 versus CD73 expression in CD4.H and CD4.I (CITE-seq; log1p(CP10K)); color indicates cell density.

We then asked whether increased functional commitment from mixed to polarized clusters was accompanied by additional molecular changes. The heatmap in **Fig. 4l** revealed shared features that collectively distinguish the poles from the mixed clusters. First, poles showed higher expression of *Klrg1* (notably in Th1 and Th2 poles), reminiscent of KLRG1 terminally differentiated effector CD8 T cells (CD8.H cluster^56^). Second, several transcription factors associated with CD4 activation were enriched (*Klf2*, *Bhlhe40*^57^), perhaps cooperating with canonical Th transcription factors to stabilize the polarized states.

Conversely, we asked whether clusters G and Q shared gene programs independent of their Th bias. Their main differences mapped to tissue migration programs: G was enriched for *Klf2*, *S1pr1*, and *Ccr7*, consistent with recirculation and lymphoid trafficking^58^, whereas Q lacked *S1pr1* and *Klf2*, the master controller of tissue exit, and expressed more *Cxcr6*, *Cxcr3*, and *Rgs1*, associated with tissue residence^59,60^ (**Fig. 4m**). Thus, the mixed states coupled graded Th program engagement with a migration dichotomy, cluster G as more migratory and cluster Q as more tissue-resident.

Several other CD4+ T cell-types have been suggested, whose molecular determinism was not always clear, such as Tr1^61^, Th9^22^, Th22^62,63^ cells. Although deeper analysis would be required for a formal conclusion, and acknowledging that very minor cell clusters might be difficult to identify, these states were not readily apparent. Concerning Tr1, *Il10* expression was distributed across many clusters, thus difficult to reconcile with the notion of a discrete subset responsible for IL10 production (**Fig. 3e**). Concerning Th22, *Il22* expression was mostly found in Th17 poles, largely parallel to that of *Il17a*, consistent with an earlier conclusion that IL-22 is not confined to a dedicated Th22 subset^64^. On the other hand, gene program (GP) analysis in a companion manuscript identified an IL-22-specific program distinct from the IL-17-associated program, consistent with partially independent regulation^65^ [**immgenT-GP]**. *Il9* was not detected among CD4+ T cells in the immgenT dataset, making it difficult to conclude about Th9 cells.

### CD4.H/I: convergence of Tfh and chronically stimulated CD4+ T cells

Mapping away from the Th2/17 poles were the two neighboring clusters CD4.H and CD4.I (**Fig. 3a**). Both shared high expression of *Izumo1r, Cd200, Eea1, Hif1a, Tox,* and reduced expression of *Cxcr6, Klf2, Sell,* (compared to all other activated CD4 clusters) (**Fig. 5a**). *Bcl6*, encoding the canonical transcription factor associated with Tfh differentiation^22,49,50^ (but not only^66–68^, was identified in both clusters albeit highest in H (**Fig. 1d**). Accordingly, these clusters were greatly reduced in T cells recognizing the I-Ab:GP66 complex in mice challenged with LCMV but lacking *Bcl6* in CD4+ T cells^69^ (**Fig. 5b)**; Finally, TCR analysis revealed preferential, albeit not exclusive, clonal sharing between I and H clusters, further substantiating the relatedness between cells in these clusters (**Fig. 3j,k**).

On the other hand, several elements argued for distinct identities. Firstly, their origin. Both clusters were represented in many samples, but CD4.H dominated in cells phenotypically defined as germinal center Tfh cells by cell surface marker (CD4⁺ CD62L^-^ CD44⁺ PD-1^high^ CXCR5^high^) after immunization with NP–OVA, a regimen known to drive Tfh formation (**Fig. 5c**). In contrast, samples enriched proportionally in CD4.I originated from tumors and chronic LCMV infection (**Fig. 5d, Extended Data Fig. 2b**), suggesting that CD4.I cells represent chronically activated T cells under conditions of persistent antigen exposure (although they are also present in some early activations, such as the flu response). However, samples exhibiting CD4.I dominance also contributed to CD4.H, and vice versa, indicating some cellular overlap rather than strict mutual exclusivity^70^, consistent with clonotype sharing (**Fig. 5e**).

Beyond their uniquely high expression of *Il21* noted earlier (**Fig. 3e**), direct comparison between CD4.H and CD4.I showed relatively few but functionally relevant differences (**Fig. 5f**). For example, CD4.H cells preferentially expressed *Cxcr5, Icos, Tox2,* and *Prdm1* transcripts (**Fig. 5f**), translating to distinctive cell surface expression of CXCR5 and PD1 in the CITE-seq data (**Fig. 5g**) This signature aligns with previous studies that defined Tfh populations^67,71,72^, suggesting that CD4.H is the main location of these GC-Tfh cells. Conversely, CD4.I diverged from CD4.H, lacking some of their key features, including *Il4* expression (**Fig. 3e**), and instead had elevated expression of *Lag3, Tnfrsf4, Itgb1,* and *Bhlhe40* (**Fig. 5f**). CD4.H and I differed diametrically in the balance of *Id2* and *Id3* expression; these E-protein interactors generally enforce quiescence vs effector differentiation, and Id2 is known to inhibit Tfh differentiation^73^. CD4.I cells co-expressed high levels of CD73 and FR4 (**Fig. 5h**), markers historically associated with T cell anergy^74–76^. Together with their dominant origin (tumors, chronic stimulation), these characteristics suggest that CD4.I cells may correspond to classically defined chronically stimulated or anergic cells.

Why do categories of T cells with such radically different roles and functions share a core transcriptional and surface program? The similarity is even more surprising since many Tfh cells occupy the specialized environment of the germinal center^77,78^, while chronically stimulated T cells, whose residence is incompletely known, are certainly outside GCs. It is important to note that both clusters appeared functionally active. Even if CD4.I cells do not dominate the cytokine production landscape, they are not inert either (**Fig. 3e**). This similarity, highlighted by co-expression of CD73 and FR4 markers, contrasts with the traditional view that these markers necessarily annotate dysfunction and instead aligns with the emerging view that CD4⁺ T cells can adopt specialized transcriptional and metabolic programs that preserve functionality despite chronic stimulation^79^. Alternatively, given that Tfh cells constantly engage pMHCII on GC B cells during the cellular ballet that drives immunoglobulin affinity maturation^77^, sustained, low-intensity antigen presentation may cause Tfh cells to exhibit characteristics of anergic cells. Although high *Bcl6* expression canonically enforces Tfh cell fate, as seen in CD4.H, its lower expression in CD4.I cells raises the possibility for non-canonical functions in sustaining chronically stimulated cells, for example, via metabolic reprogramming or adaptation^67,80^. Together, these data reveal a striking convergence between Tfh and chronically stimulated CD4+ T cells, despite arising from distinct immunological contexts.

### CD4+ memory T cells landscape

Defining the molecular programs of CD4 memory T cells has been challenging. There are technical limitations (e.g., antigen-specific CD4+ T cells at memory time points contract to very low numbers). It is unclear whether the framework of memory CD8 T cells can be applied to CD4+ T cells^7,81–83^. As a result, descriptions of CD4 memory have become complex, combining CD8-derived classifications (e.g., MPEC/SLEC, Tcm, Tem, Trm) with those of helper CD4+ T cell programs. We asked how memory CD4+ T cells are distributed within the immgenT framework.

We operationally defined memory CD4+ T cells as antigen-specific cells present after resolution of infection or allergen exposure, at a time when antigens loaded on APCs would be expected to have cleared, a key criterion in defining memory^84^. CD4+ T cells specific for the 2W antigen of engineered *Listeria monocytogenes* were profiled over a fine-grained time-course (**Fig. 6a**). This strain of *Listeria* is typically cleared by day (d) 10 (post-infection, pi). During the effector phase (d6-d10), clusters G, H and I predominated (**Fig. 6b**), with a smaller representation of the Th1-like R polarized cluster. Through the transition and into memory time points, this distribution largely persisted, albeit with a decrease in cells in CD4.R and the appearance of cells in the resting clusters A and C (∼15% altogether, these memory cells maintained some *Cd44* expression, **Extended Data Fig. 1a**). The bias in differentiated programs in effector phase cells (**Fig. 6c**) was also conserved at memory-phase (**Fig. 6d**): Th2- and Th17-like programs defined above (**Fig. 4a-d**) remained extinct, while Th1- and Tfh-like programs persisted (**Fig. 6c,d**, **Extended Data Fig. 6a,b**). In a second set, CD4+ T cells from the Smarta TCR transgenic mouse, specific for LCMV GP66:I-Aᵇ, were transferred into a normal host prior to infection with LCMV-Arm (**Extended Data Fig. 6c,d**). During the effector phase (d7 pi), clusters G, H and R predominated, consistent with Th1-like and Tfh-like responses to LCMV^41,83,85–87^ (**Extended Data Fig. 6e**). Here again, these clusters persisted in the memory phase and the deviation towards Th1 and Tfh programs was also maintained (**Extended Data Fig. 6f**). Thus, in the systems analyzed here, most antigen-specific memory CD4+ T cells map to the mixed clusters, but they retain the programmatic bias of the effector phase.

**Figure 6.**
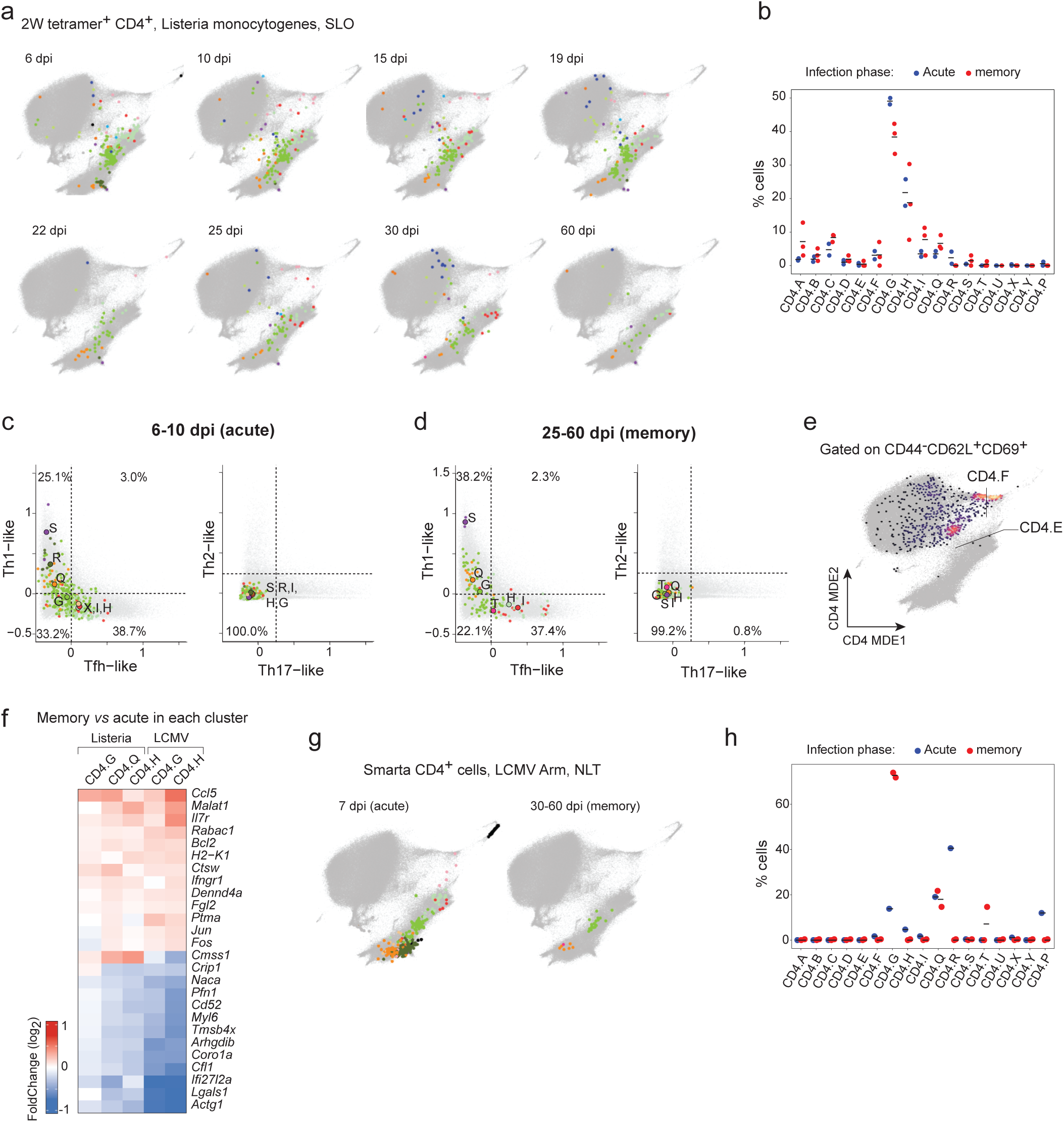
Memory CD4+ T cells map to the same clusters as effector cells. **(a)** CD4 MDE showing the cluster distribution of *Listeria*-specific CD4+ T cells in secondary lymphoid organs (SLO) across eight time points post-infection (6–60 dpi) (IGT43). 2W tet⁺ denotes antigen-specific cells recognizing *Listeria monocytogenes* engineered to express the 2W peptide. Cells colored by cluster identity; background immgenT CD4+ T cells in grey. **(b)** Quantification from (a) showing the proportion of CD4 clusters at the acute phase (6–10 dpi, blue) versus memory phase (25–60 dpi, red). Each dot represents one sample; bars indicate the mean. **(c,d)** Scatterplots showing pairwise expression of Th programs in *Listeria*-specific CD4+ T cells from (a) at acute and memory phases. Larger labeled circles indicate the mean program expression of each activated cluster (G–Y). **(e)** Heatmap of genes differentially regulated between memory and acute phases, shared across clusters, in both *Listeria* infection and LCMV Armstrong infection in SLO. **(f)** CD4 MDE showing the cluster distribution of LCMV-specific CD4+ T cells (SMARTA) in non-lymphoid tissues (small intestine, prostate, salivary gland, bone marrow, lung) at 7 dpi (acute phase, left) and 30–60 dpi (memory phase, right) (IGT36,38,40). Cells are colored by cluster identity; background immgenT CD4+ T cells in grey. **(g)** Quantification from (f) showing the proportion of CD4 clusters at acute (7 dpi, blue) versus memory (30–60 dpi, red) phases in NLT. Each dot represents one sample; bars indicate the mean.

We expanded the analysis with published datasets mapped to the immgenT CD4 reference using T-RBI. 2W-tetramer–positive cells at different time points after *Listeria* infection from Osum et al.^60^ were consistent with the present results, with perhaps a stronger representation of the resting clusters (**Extended Data Fig. 6g,h**). CD4+ T cells expressing the MHC-II-restricted Smarta TCR during LCMV Armstrong infection^69^ again showed memory cells mapping to the same clusters as effector cells, with a preponderance of mixed clusters G and Q and a decrease in polarized cells (**Extended Data Fig. 6i,j**). Memory response to malaria^88^ also showed a similar cluster distribution **(Extended Data Fig. 6k).**

However, despite these similarities, cells assigned to each cluster did not completely overlap between the effector and memory phases. These differences were brought out by a cluster-by-cluster comparison of gene expression in acute and memory cells in LCMV-specific Smarta and Listeria-specific 2W CD4+ T cells (**Fig. 6e**). Memory/effector differences were conserved across all three clusters tested. This “memory signature” included the overexpression of a few evocative genes, such as *Il7r* and *Bcl2* which are central to T cell survival and homeostasis^60,89^, and a few others whose role in memory maintenance is less expected (*Malat1*^90^, *Ccl5*^91^, or *Ctsw)*.

Finally, we examined whether the cluster distribution observed in SLOs was conserved in non-lymphoid tissues (NLTs). In the same experiments as described above, Smarta T cells were recovered from several NLTs (lung, submandibular gland, bone marrow, prostate, and small-intestinal lamina propria (SILP) (**Fig. 5f,g**). During the effector phase, the cluster distribution in NLTs was broadly similar to that observed in SLOs (**Fig. 5b, Extended Data Fig. 6l**), but with an over-representation of the cluster Q and R. At memory time points, the same “retreat to the mixed clusters” was observed as in SLOs, cluster R cells disappearing and replaced by clusters G and Q (with slightly more cluster Q than in SLOs).

In summary, memory CD4 states exhibited four key features: (i) no particular location in the phenotypic space for memory CD4+ T cells (ii) these cells remained in the same clusters as effector cells, but with a preponderance of “mixed clusters” cells and disappearance of the polarized phenotypes; (iii) re-emergence, more marked in some models, of a substantial quiescent population (clusters A-C); and (iv) a subtle “memory signature” shared across clusters, that may explain long-term homeostatic maintenance of the CD4+ memory pool. These features contrast with CD8 memory, in which effector and memory cells segregate more clearly into distinct clusters (**Extended Data Fig. 6m** and accompanying immgenT-CD8 manuscripts^56^).

### Heterogeneity in the resting pool

A large compartment of CD62L⁺CD44⁻ CD4+ T cells (clusters A–F; **Fig. 1c**) corresponded to the well-described “resting” populations, enriched in typical quiescence-associated transcripts (*Sell, Lef1, Satb1, Bach2, Tcf7, Klf2, Ccr7*) and reduced expression of effector genes relative to activated clusters (G–Y) (**Extended Data Fig. 7a**). They expressed high levels of CD45RB, CD55, and LY6C surface proteins (**Fig. 7a**). However, this compartment was not uniform and instead resolved into six robustly distinct clusters, in accordance with a few studies that recently documented heterogeneity in the resting/naïve pool of CD4+ T cells^92–95^ (**Extended Data Fig. 7b-c**). Clusters A and C were the most abundant populations, enriched in SLOs (**Fig. 7b**). Cluster B was notable for its transcriptional signature enriched *Il7r, Klf2, Ccr7*. Previous work indicated that this cluster is restrained by VISTA and expands in its absence^92^. Here, this cluster was particularly enriched in NLTs (**Fig. 7b,c**) and uniquely expressed *Ifngr1*, possibly reflecting a poised state of activation in tissues. Clusters D–F were smaller (<5% of cells) but exhibited distinct activation-associated programs (**Fig. 7d, Extended Data Fig. 7c**). Cluster D was marked by a high level of Type-I IFN signature genes (e.g., *Isg15, Stat1, Irf7, Ifit1*), consistent with reports of small T cell clusters uniquely exposed (or responsive to) IFN^28,96,97^. Clusters E and F expressed CD69 (**Fig. 7e)** and increased CD5, along with TCR-induced genes (*Nr4a1, Egr1, Egr2*) (**Fig. 7d)**, consistent with TCR tuning. Consistent with their resting phenotype, clusters A–E displayed highly diverse TCR repertoires with minimal clonal expansion, and little clonal overlap with activated clusters (**Fig. 3i–k, 7f**). In contrast, cluster F showed modest clonal expansion during immune challenge and shared clonotypes with activated populations, suggesting a possible intermediate state towards activation. Compared with antigen-driven clonal responses, systemic bystander effects of immune perturbation were more evident in the resting populations of spleen and lymph nodes (**Fig. 7g**).

**Figure 7.**
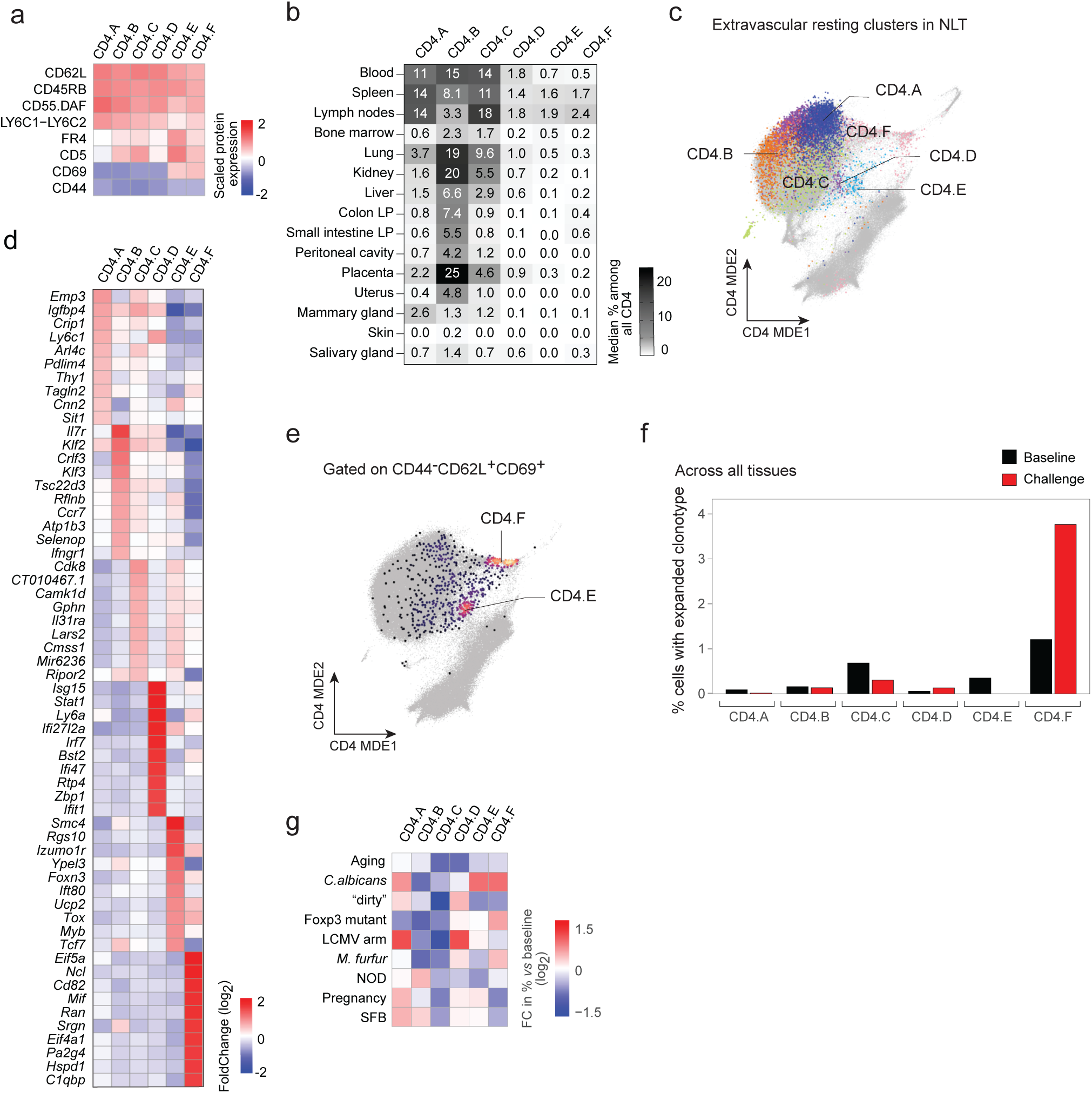
Resting CD4+ T cells heterogeneity. **(a)** Heatmap showing expression of the most discriminative surface markers across CD62L⁺ CD44⁻ resting clusters (CD4.A–F), measured by CITE-seq (Z-score). **(b)** Heatmap showing the median proportion of resting clusters across tissues at baseline. **(c)** CD4 MDE showing extravascular cells mapping to resting clusters in non-lymphoid tissues (lung, small intestine, prostate, submandibular gland, and skin), based on in vivo labeling of intravascular cells by intravenous anti-CD45 antibody prior to tissue harvest. Cells are colored by cluster identity. **(d)** Heatmap showing the top 10 genes upregulated in each cluster compared to all other resting clusters (adjusted *P* < 0.05). **(e)** CD4 MDE showing CD44⁻ CD62L⁺ CD69⁺ cells identified by bioinformatic gating using CITE-seq. **(f)** Bar plots displaying the percentage of cells with expanded clonotypes (≥2 cells per clonotype) in each resting cluster (CD4.A–F) at baseline (black) and during immune challenge (red). **(g)** Heatmap showing changes in the proportions of resting CD4⁺ T cell clusters in secondary lymphoid organs under different immune conditions. Color intensity indicates the log2 fold change relative to baseline.

Overall, the “resting continent” of CD62L⁺CD44⁻ CD4+ T cells was heterogeneous and did not represent a uniform SLO-restricted resting state. Instead, this compartment included subsets enriched in NLTs and populations exhibiting early activation and clonal expansion programs.

### immgenT CD4 flow cytometry panel

Leveraging the 128 surface markers measured by CITE-seq, we defined a reduced 12-marker flow cytometry panel to resolve the CD4+ T cell landscape (**Extended Data Table 5**). While many markers have been previously described^7,60,83^, immgenT enabled systematic evaluation of whether their combinations mapped to discrete regions of the MDE or instead spanned multiple states.

The foundational markers were CD62L and CD44 (**Fig. 8a-e**). CD62L⁺ CD44⁻ cells mapped to resting clusters A–F, whereas CD62L⁺ CD69⁺ cells corresponded to clusters E and F (**Fig. 8a,b**; **Fig. 7**). Within the CD44⁺ compartment, CXCR6 expression defined a secondary axis separating clusters G–I from Q–Y (**Fig. 8a,b**). Among CXCR6⁻ cells (**Fig. 8d**), CXCR5hi PD-1hi cells mapped to CD4.H, consistent with a GC-Tfh phenotype (**Fig. 8d**), whereas CXCR5lo PD-1hi cells mapped to CD4.I (**Fig. 8d**). CXCR5lo PD-1lo/neg cells corresponded to the mixed state G (**Fig. 8d**). As discussed in the previous sections, other canonical gating strategies converged on cluster G, including CD44⁺ CCR7⁺ (**Extended Data Fig. 1b**) and CD44⁺ CD62L⁺ cells (**Extended Data Fig. 8a**), commonly used to define central memory-like populations. Similarly, CD73hi FR4hi cells, associated with anergy, mapped broadly to clusters G–I (**Fig. 5i**; **Fig. 8a,b**).

**Figure 8.**
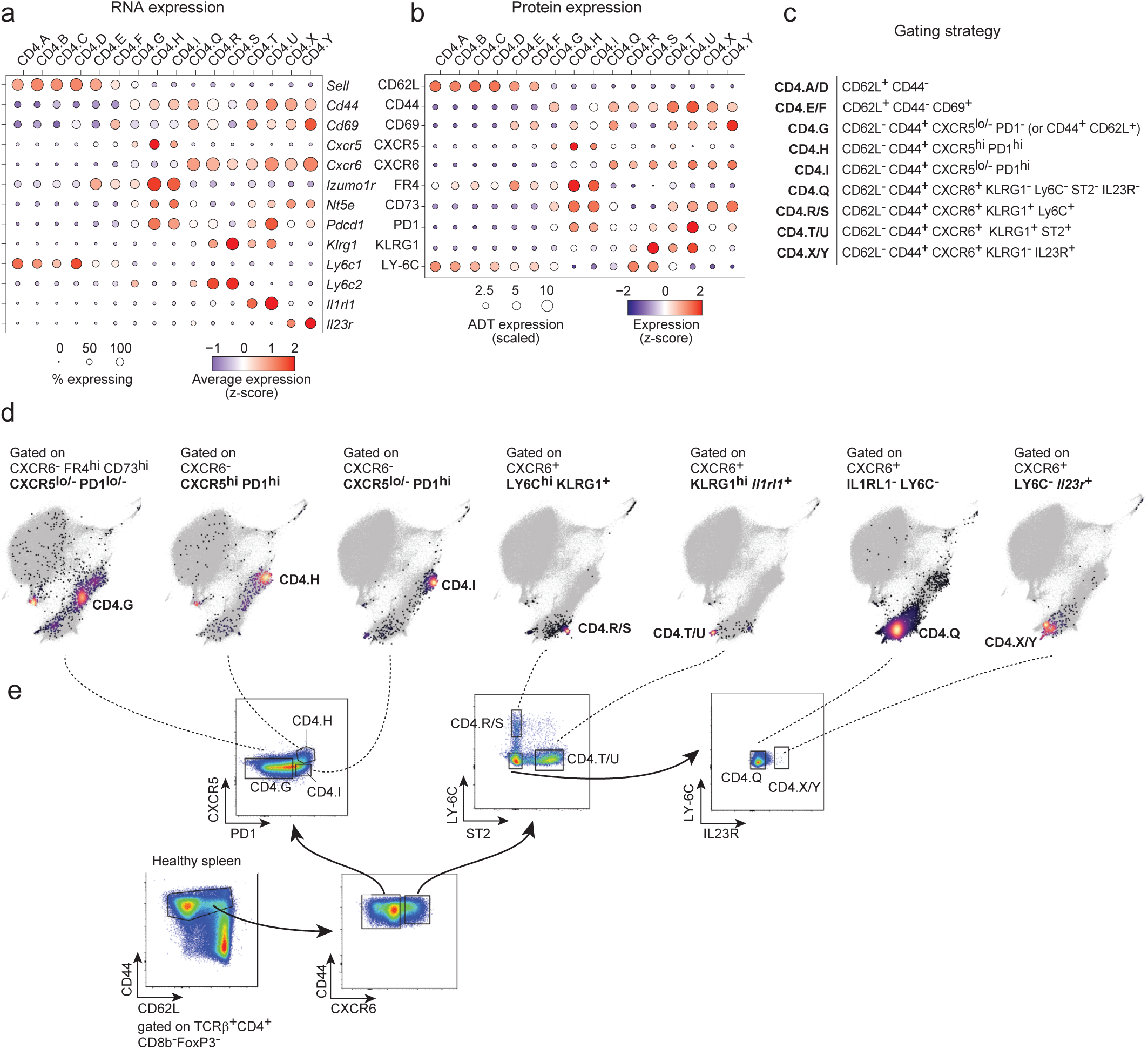
immgenT CD4 panel: a flow cytometry strategy to resolve the CD4+ T cell landscape. **(a,b)** Dot plots of the immgenT CD4 surface marker panel showing gene (a) and protein (b) expression across clusters. Color indicates scaled expression (Z-score); dot size indicates the fraction of expressing cells. Of note, ST2 and IL-23R were not robustly detected by CITE-seq, and chemokine receptors such as CXCR5 and CXCR6 showed weak signal (see Methods). **(c)** Table summarizing the main gating strategy for each cluster (complete strategy in **Extended Data Table 5**). **(d)** CD4 MDE highlighting cells bioinformatically gated using CITE-seq and the strategies described in (c). Cells pre-gated on CD44⁺ CD62L⁻ in addition to the gates shown above each plot. Color indicates cell density; background immgenT CD4+ T cells in grey. **(e)** Representative flow cytometry plots illustrating the gating strategy used to identify CD4+ T cell clusters. Data shown from spleen at baseline (C57BL/6J mouse).

In an opposite region of the MDE, CXCR6+ cells encompassed clusters Q–Y (**Fig. 8a,b**). Th1 (R/S) and Th2 (T/U) polarized clusters were marked by KLRG1hi expression (**Extended Data Fig. 8b**) and further distinguished by Ly6C and ST2, respectively (**Fig. 8d,e**). Th17-like (XY) cells were identified by IL-23Rhi expression (**Fig. 8d,e**). CXCR6⁺ cells lacking KLRG1, Ly6C, IL-23R, or ST2 mapped to the mixed Q state, consistent with reduced helper T cell effector programs (**Fig. 8d,e**). **Fig. 7e** and **Extended Data Fig. 8c-f** illustrate applications of this panel to splenic CD4⁺ T cells as well as to lung and colon samples. Intracellular transcription factor staining further supported assignment of the polarized clusters to specific Th programs, with T-bet, GATA3, and RORγt enriched in Ly6C⁺ Q/RS, ST2⁺ TU, and IL-23R⁺ XY gated populations, respectively (**Extended Data Fig. 8g-i**).

Overall, this 11-marker panel captured the major axes of CD4+ T cell heterogeneity and recapitulated transcriptionally defined states across tissues and conditions. Beyond this panel, the Rosetta web application (rosetta.immgen.org) enables flexible evaluation of all 128 markers and their mapping onto the immgenT MDE, allowing assessment of whether they resolve discrete regions of the embedding or span multiple states.

## Discussion

By combining datasets from an unprecedented number of tissues and conditions, immgenT distilled the diversity of CD4⁺ T cell states into 20 robust and reproducible clusters that capture resting and antigen-experienced compartments, unifying states arising from diverse challenges and locations. These revealed a graded polarization model of CD4⁺ T cell differentiation, in which a minority of cells occupy highly polarized states, coexisting with a broader pool of activated but less committed cells that populate intermediate states in which the same Th programs are present, but less sharply and non-exclusively deployed. The data further highlight substantial overlap between Tfh-like states and those that arise under chronic antigen exposure and show that memory cells largely retain phenotypic configurations similar to those of their effector counterparts.

These observations enable the definition of core *in vivo* Th signatures that are broadly applicable and not biased by simple pairwise comparisons, which often do not generalize. They support a model of CD4+ T cell heterogeneity structured by layered gene programs, in which variation arises from combinations and levels of activity of these programs, rather than from an array of discrete cell states. This framework reconciles the models of CD4+ T cell differentiation that evolved over time. Classical models emphasized terminally differentiated Th states (Th1/Th2/Th17/Tfh) with mutually exclusive programs^13,15,16,21,50,98–100^. More recent models have emphasized the notion of continua in differentiated CD4+ T cells, rather than discrete entities, with graded phenotypes rather than binary fate decisions^22–25,27,28,94,101^. We propose that highly differentiated Th cells with exclusive programs in the polarized clusters represent *in vivo* equivalents of the canonical states defined *in vitro*, while the continua of biased cells with lower program activity and exclusivity map to intermediate mixed states. The mechanisms driving the emergence of the poles from mixed states remain unclear. Signals delivered via the TCR^102^ or biasing cytokines (e.g., IL-12)^103^, interactions with APCs^104^, may trigger the self-reinforcing loops based on cytokines or TFs (T-bet, Blimp-1, Bcl6, GATA3, RORγt, IRF4^105^), and ZEB2^106^ or KLF2^107^ may play a role in narrowing or broadening the programs. The TCR data further informed the relationships between these states: sharing of TCR clonotypes was most prevalent between differentiated cells from related clusters (T and U, X and Y), but was nevertheless present between more distant ones. This could result from plasticity and programmatic switching, or because descendant cells of a single naïve cell quickly diverged. Either way, there is no obligate relationship between TCR clonotype and cell fate during an immune response.

This organization encompasses several longstanding observations. First, large fractions of activated CD4+ T cells that do not produce canonical cytokines have long been observed by flow cytometry (e.g., ref.^108^). Rather than representing alternative polarized states, these activated but non-producing cells mapped here to the mixed clusters G and Q. Second, polyfunctional cells co-expressing cytokines (most often IL-17 and IFN-γ) have been repeatedly documented^15,21–25^, which is readily explained by the combinatorial expression model in clusters G and Q. Similarly, there are reports of Tfh-like cells with elements of Th1/2/17 signatures^5,7^. In the immgenT framework, these cells position within the mixed cluster G. The presence of CD4.G cells in non-lymphoid tissues further aligns with descriptions of resident Tfh-like populations^109,110^. Thus, cluster G may unify diverse nomenclatures (Tfh-like, M-Tfh, Th0, Tfh-Trm, Tfh1, Tfh2, Tfh17, Tcm-like) that have been proposed^5,60,66,109–112^.

More generally, if we define the polarized clusters as the archetypal Th states, how do we account for the subsets of Th cell-types that have been described? For instance, the distinction between pathogenic vs non-pathogenic Th17 cell^14,26^ has important functional correlates, but there is no clear consensus on a corresponding transcriptional signature, beyond pathogenic Th17 populations being more prone to express IFNγ. Rather than distinct subsets, Th17 cells of varying pathogenicity may simply correspond to the fluidity within the mixed CD4.Q. Similarly, it is well recognized that the Th1 concept actually covers several entities which have the *Tbx21/Ifng* program in common. The immgenT data map this diversity, spanning clusters R and S and extending into Q and G, present in multiple infection and tumor models . CD4.S cells resemble intravascular CX3CR1⁺ CXCR3⁻ cytotoxic cells^113,114^, whereas a CX3CR1⁻ CXCR3⁺ population likely corresponds to cluster R^60,83,106,113,114^. Th1-like cells in Q are consistent with tissue-resident states^60^, while cluster G cells display less differentiated or central-memory–like Th1 features^106^.

These findings highlight the limitations of nomenclatures that have been derived over the past decades, from a mix of criteria (functional (“Tfh”), antigen exposure (“CD4act”) or residence (“CD4.Trm”). Cells of a given label may map to a single cluster, but the reverse isn’t the rule: in most cases, each cluster encompasses multiple previously described populations. Accordingly, and as a general principle emerging from immgenT, individual clusters cannot be fully annotated using existing classification schemes. **Extended Data 6** attempts to summarize the correspondence between immgenT clusters and previous descriptors (this table may evolve with community input). We therefore propose using the immgenT framework as a shared reference (using the T-RBI integrator to map scRNAseq data, or the marker panel of Fig.7 to integrate flow cytometry data).

Finally, our analysis shows that CD4+ T cells may undergo fewer transcriptional changes between peak clonal expansion and later memory cell emergence than CD8+ T cells. Distinct clusters of short-lived effector cells and memory precursor effector cells that are apparent for CD8+ T cells are not present in the CD4+ T cell dataset^56^. The transcriptional changes that determine the CD4+ T cell clusters, for example, the Th1/Th2/Th17 and tissue residence/recirculation programs occur early and are retained by the memory cells that survive the contraction phase along with subtle changes that foster cell longevity and quiescence^115^.

In conclusion, the immgenT framework provides a reference to anchor CD4+ T cell heterogeneity for future investigation. The presence of each cluster across diverse biological contexts suggests that these states may represent attractor/shared states that can be reached through multiple differentiation trajectories.

## Supporting information

Extended Data Table 1 - Sample Metadata

Extended Data Table 2 - Cluster Signatures

Extended Data Table 3 - TCR Clonotypes

Extended Data Table 4 - In vivo Helper T cell signatures

Extended Data Table 5 - CD4 Flow Cytometry Panel

Extended Data Table 6 - immgenT Previous Nomenclature

## Acknowlegments

We thank the many colleagues who were consulted at various stages of this project. This work was funded by a grant from the NIH to the ImmGen consortium (R24-072073).

## Author Contributions

NM, DB, AL, GCWB, RL, TS, KCO, AF, TH performed the experiments; NM, ZZ, DB, AL, GCWB, RL, TS, LN, DZ participated in the computational processing and analyses; DZ, JM, MKJ, CB, KC, MS, SB, SS, MP and immgenT principal investigators designed the study, provided complementary funding, and oversaw the experiments; DZ, CB, MKJ and JM wrote the manuscript with input from other authors.

## Collaborators

Participants in the immgenT Project include:

Aaron Liu^1^, Alexander Chervonsky^2^, Alexandra Cassano^2^, Alia Welsh^3^, Amir Ferry^11^, Ananda Goldrath^11^, Andrea Lebron-Figueroa^5^, Ankit Malik^2^, Anna-Maria Globig^4^, Antoine Freuchet^2^, Bana Jabri^2^, Charlotte Imianowski^6^, Christophe Benoist^5^, Claire Thefaine^7^, Dan Kaplan^6^, Dania Mallah^5^, Dario Vignali^6^, David Sinclair^5^, David Zemmour^2^, Derek Bangs^8^, Domenic Abbondanza^2^, Enxhi Ferraj^9^, Eric Weiss^6^, Erin Lucas^7^, Evelyn Chang^9^, Gavyn Bee^10^, Giovanni Galletti^11^, Ian Magill^5^, Iliyan D Iliev^12^, Jinseok Park^5^, Joonsoo Kang^9^, Jordan Voisine^2^, Josh Choi^5^, Julia Merkenschlager^13^, Jun R. Huh^5^, Katharine Block^7^, Ken Cadwell^10^, Kennidy K. Takehara^11^, Kevin Osum^7^, Laurent Brossay^14^, Laurent Gapin^15^, Liang Yang^5^, Lizzie Garcia-Rivera^1^, Marc K. Jenkins^7^, Maria Brbic^16^, Maria-Luisa Alegre^2^, Marion Pepper^8^, Mariya London^17^, Matthew Stephens^2^, Maurizio Fiusco^16^, Melanie Vacchio^3^, Michael Starnbach^5^, Michel Nussenzweig^13^, Mitch Kronenberg^18^, Myriam Croze^19^, Nalat Siwapornchai^5^, Nathan Morris^12^, Nicole E. Scharping^11^, Nika Abdollahi^19^, Nitya Mehrotra^2^, Odhran Casey^5^, Olga Barreiro del Rio^5^, Paul Thomas^20^, Peter Carbonetto^2^, Remy Bosselut^3^, Rocky Lai^9^, Sam Behar^9^, Sam Borys^14^, Sara E. Hamilton^7^, Sara Mostafavi^8^, Sara Quon^11^, Serge Candéias^21^, Shanelle Reilly^14^, Shanshan Zhang^5^, Siba Smarak Panigrahi^16^, Sofia Kossida^19^, Stefan Muljo^3^, Stefan Schattgen^20^, Stefani Spranger^22^, Steve Jameson^7^, Susan M. Kaech^1^, Takato Kusakabe^12^, Taylor Heim^22^, Tianze Wang^8^, Tomoyo Shinkawa^9^, Ulrich von Andrian^5^, Val Piekarsa^5^, Véronique Giudicelli^19^, Vijay Kuchroo^5^, Woan-Yu Lin^12^, Ziang Zhang^2^

1. NOMIS Center, Salk Institute for Biological Sciences, 2. The University of Chicago, 3. National Institutes of Health, 4. Allen Institute for Immunology, 5. Harvard Medical School, 6. Dept of Dermatology and Immunology, University of Pittsburgh, 7. University of Minnesota, 8. University of Washington, 9. UMass Chan Medical School, 10. University of Pennsylvania, 11. University of California San Diego, 12. Weill Cornell Medicine, 13. The Rockefeller University, 14. Brown University, 15. University of Colorado Anschutz Medical Campus, 16. Swiss Federal Institute of Technology, Lausanne, 17. New York University, 18. La Jolla Institute, 19. IMGT, Université de Montpellier, 20. St. Jude Children’s Research Hospital, 21. Alternative Energies and Atomic Energy Commission, Grenoble, 22. Massachusetts Institute of Technology

## Methods

### Mice

Mice used in the immgenT dataset are described in detail in the immgenT Cosmology manuscript (**Extended Data Table 1**) and on the immgenT website (https://immgen.org/ImmGenT/), including sex, age, and genetic background. With rare exceptions, experiments were performed using C57BL/6 (B6) mice, most of which were sourced from The Jackson Laboratory. Experimental conditions, including infection models, immunization strategies, and tissue processing protocols, are detailed in the immgenT Cosmology manuscript (**Extended Data Table 1**), the GEO GSE297097 dataset, and the immgenT website. Both male and female mice were used.

The flow cytometry experiments presented in the present study were conducted on mice bred and maintained under specific pathogen–free conditions at the University of Chicago, in accordance with procedures reviewed and approved by the University of Chicago Institutional Animal Care and Use Committee (IACUC) guidelines (Protocol ID: 72714).

### ImmgenT dataset: experiments and data processing

The immgenT dataset comprises 66 experiments of single-cell RNA-seq, CITE-seq (128-plex), and paired TCRαβ sequencing (10x Genomics 5′ v2 platform), corresponding to 80 encapsulation runs. Each encapsulation is assigned a unique “IGTx” identifier (IGT1–96) used to track datasets. In a minority of cases, two IGT identifiers denote parallel encapsulations of the same biological samples. Individual cells are indexed by a unique IGT.cellID, and samples are tracked using hashtag identifiers (IGT.HT).

A detailed description of data acquisition and processing is provided in the immgenT Cosmology manuscript^29^. Briefly:

*Logistics*. Experiments were conducted across multiple laboratories in the United States. Participating laboratories carried out mouse treatments (e.g., infection or immunization) and tissue processing with their established materials and reagents. The immgenT research assistant (IM) traveled to the site, helped with sample CITE-seq labeling and cell sorting, and performed encapsulation and library preparation. Each experiment typically included ∼10 hashtagged and pooled samples, and a standardized spleen control for batch-effect assessment. Library construction and sequencing were centralized at the Broad Institute.

*Sample processing, library construction, and sequencing.* Flow cytometry–sorted T cells were multiplexed using TotalSeq-C hashtag^116^ antibodies (BioLegend), enabling pooling prior to encapsulation. Cells were stained with a custom 128-antibody TotalSeq-C panel and subjected to joint single-cell RNA, surface protein (CITE-seq), and TCR sequencing using the 10x Genomics 5′ v2 platform. Libraries were sequenced on an Illumina NovaSeq.

*Data processing and quality controls.* Gene expression, protein, hashtag, and TCR count matrices were generated using Cell Ranger, and hashtag-based demultiplexing was performed in Seurat^117,118^. Cells were filtered based on RNA and protein quality-control metrics detailed in the immgenT Cosmology manuscript - Extended Data Note 1.

*Data integration.* Datasets were integrated using totalVI^30,119^, with the 10x Genomics lane (IGT) specified as a batch covariate. The model was trained on all detected genes and proteins using a 30-dimensional latent space (default parameters).

*Dimensionality reduction.* Dimensionality reduction was performed using Minimum Distortion Embedding (MDE) implemented in the pyMDE^120^ library via the pymde.preserve_neighbors() function with default parameters. In contrast to UMAP, which is stochastic and graph-based, pyMDE preserves both local neighborhood structure and global geometry with minimal distortion. After computing the embedding on the full immgenT dataset (IGT1–96), coordinates were reused to anchor immgenT cells as a reference, enabling projection of new datasets without altering the original embedding. This enabled the construction of shared T cell reference embeddings (both all-T and lineage-specific, including CD4-specific embeddings).

*Cell clustering and annotation.* Clustering was performed in the totalVI latent space using the Louvain algorithm (Seurat FindClusters). Clustering solutions were evaluated across resolutions (0.5–4) and optimized using silhouette scores to balance over- and under-clustering. Clusters were merged or split based on consistency across samples and coherence of RNA and protein expression profiles (see immgenT Cosmology manuscript, Extended Data Note 3). In total, 1,710 of 719,580 cells (0.02%) could not be confidently annotated and were labeled as “unclear” and excluded. Small provisional clusters (<1% of cells) are denoted with a “w” prefix.

CD4 T cells were identified as CD4⁺Foxp3⁻ conventional αβ T cells based on combined RNA and protein expression of canonical markers and TFs, including *Cd3e, Trbc1, Trbc2, Trgc1, Trdc, Cd4, Cd8a, Cd8b1, Foxp3* and *Zbtb16*, as well as surface proteins CD3, TCRβ, THY1.2, CD4, CD8A, CD8B, and TCRγδ. Other TCRαβ⁺ CD4⁺ populations, including Zbtb16⁺ T cells (Tz; iNKT and MAIT cells) and CD4⁺Foxp3⁺ Tregs, formed distinct clusters and were excluded from this definition.

*CITE-seq analysis.* The full panel of 128 antibodies and associated quality metrics are provided in **Extended Data Table 4** and **Extended Data Note 1** of the immgenT Cosmology manuscript^29^. Antibody performance was evaluated using RNA–protein correlation and protein dynamic range, stratifying antibodies into high-, intermediate-, and low-performance groups (n = 61, 41, and 26, respectively). Overall, most of the CITE-seq panel performed robustly, enabling dropout-resistant protein quantification that closely parallels flow cytometry and supports both high-dimensional analyses and conventional gating strategies. For flow cytometry–like plots, protein counts were normalized using Seurat’s LogNormalize method (log1p CP10K).

*UMAP and clustering of individual IGT datasets.* UMAP and clustering were also performed on individual IGT datasets, generating dataset-specific UMAPs used in the Rosetta2 databrowser and in some immgenT manuscripts. These used the standard Seurat functions: NormalizeData(normalization.method = "LogNormalize", scale.factor = 1e4) %>% FindVariableFeatures(selection.method = "vst", nfeatures = 2000) %>% ScaleData(features = VariableFeatures(.)) %>% RunPCA(features = VariableFeatures(.), npcs = 50) %>% FindNeighbors(dims = 1:30) %>% FindClusters(resolution = 1).

### immgenT Reference-Based Integration (T-RBI)

The full methods are described in the immgenT Cosmology manuscript^29^. Integration of query datasets using T-RBI is available at https://www.immgen.org/ImmGenT/.

Reference-based integration of external datasets (T-RBI) was performed using scVI/scANVI^39,40,119^, and pyMDE^120^. Query datasets are first filtered for T cells using gene signature scoring, with γδ T cells identified and analyzed separately from αβ T cells. scVI models were trained jointly on reference (immgenT) and query data to learn a shared latent space, followed by scANVI to predict lineage and sub-lineage annotations with associated confidence scores. Cells with low-confidence predictions were iteratively reassigned until convergence. To quantify whether unannotated cells occupy regions of the embedding space not represented in the annotated reference (i.e., potential novel T cell states), we defined a discovery score based on a local k-nearest neighbor (kNN) distance ratio between query and reference cells. Scores >1 indicate that a cell is locally closer to other unannotated cells than to annotated cells, consistent with occupancy of under-represented or novel regions of the reference space, whereas scores ≤1 indicate embedding within previously annotated regions. The resulting latent representations were then used to project query cells onto the immgenT reference embeddings using pyMDE with anchored constraints, preserving the original atlas structure while incorporating new data.

Final outputs include lineage annotations, confidence scores, discovery scores, and coordinates in both global (all T MDE) and CD4-specific MDE embeddings, enabling direct comparison of query datasets within the immgenT framework.

### Differential Gene Expression

*Limma*. Differential gene expression was performed using a pseudobulk linear modeling framework based on limma^121,122^ and edgeR. Briefly, raw RNA counts from the Seurat object were aggregated by cell cluster (annotation_level2) and experiment (IGTHT), and normalized using trimmed mean of M-values (TMM). A design matrix encoding cluster–experiment combinations was constructed, and gene-wise linear models were fit to the normalized expression matrix using limma. Differential expression was assessed using empirical Bayes moderation of variance estimates, and statistical significance was determined with Benjamini–Hochberg correction for multiple testing.

*FlashierDGE*. Fast differential expression was performed in the flashier^123,124^ EBMF semi-NMF framework (described in more details in the companion immgenT-GP manuscript) by leveraging the learned gene programs (factors) and cell loadings from a 200-factor model fit to log-normalized expression. For a given comparison, mean factor loadings were computed separately for group 1 and group 2, and the difference in mean loadings (Δloadings) provided differential gene-program activity. Gene-level differential expression was then reconstructed directly from the factorization as F×Δloadings, yielding per-gene log fold changes. The same framework also returned average expression estimates for both programs and genes from the group-wise mean loadings and reconstructed group means. The function is available in ZemmourLib package (FlashierDGE).

### Quantitative analysis of Th cell programs

*In vivo Th1 (RS), Th2 (TU), Th17 (XY), and Tfh (HI) signatures* were defined as genes meeting the following criteria: minimum expression (log1p(CP10K) ≥ 0.25) in pseudobulk expression of each of the two polarized clusters, at least 1.5-fold enrichment relative to the highest-expressing alternative polarized cluster, and adjusted *P* < 10⁻⁵.

*Single-cell quantification of Th signatures.* Gene signature scores were computed using Seurat’s AddModuleScore() function.

*Th program co-expression statistical analysis.* To test whether the frequency of cells co-expressing two Th programs could be explained by the random assortment of individual program activities, we compared observed and expected co-expression frequencies under an independence model. This analysis was performed across all six pairwise combinations of Th1, Th2, Th17, and Tfh programs. For each pair (Thx, Thy), we computed the empirical joint exceedance probability P(Thx > x, Thy > y), defined as the fraction of cells simultaneously exceeding both thresholds, and compared it to the independence expectation P(Thx > x)P(Thy > y), calculated as the product of marginal probabilities. Deviation from independence was quantified as: K(x,y) = P(Thx > x, Thy > y) / [P(Thx > x) P(Thy > y)] where K < 1 indicates reduced co-expression (mutual exclusion). This ratio was evaluated over a two-dimensional grid spanning the 1st to 99th percentiles of each Th program. To further characterize mutual exclusion along a single axis, scores were transformed to empirical quantiles using the probability integral transform. Fixing qy = 0.8 (80th percentile of Thy), we computed K(qx, 0.8) across qx values ranging from 0.02 to 0.98. Statistical significance at each qx was assessed using a one-sided Fisher’s exact test (alternative: exclusion, i.e., odds ratio < 1), with p-values adjusted using the Benjamini–Hochberg procedure to control the false discovery rate (FDR < 0.05). Confidence intervals were estimated from 500 bootstrap replicates.

### TCR Clonotype sharing and overlap analysis

Clonotype sharing across CD4 T cell clusters was assessed using matched single-cell TCR sequencing data as described in the immgenT-TCR manuscript^33^. Clonotypes were defined based on paired TCRαβ and TCRβ sequences with full nucleotide-level identity in the junction sequences. For each cluster, the proportion of cells with expanded clonotypes was calculated as the fraction of cells with clonotypes observed more than once in the sample. To quantify overlap between clusters, pairwise clonotype sharing was computed using the Jaccard index on sets of unique clonotypes per cluster. For visualization, a weighted network of cluster relationships was constructed, where edge weights correspond to the Jaccard index. Edges corresponding to only one shared clonotype were omitted for clarity and the network was overlaid onto the MDE embedding using cluster centroids.

### immgenT CD4 flow cytometry panel design and testing

*Panel design.* For each lineage, COMETSC^125^ was applied to normalized protein expression matrices together with MDE coordinates and cluster annotations, allowing up to two-gene combinations (-K = 2) to identify combinatorial markers. In combination with literature curation, extended marker combinations were evaluated for CD4 cluster specificity using CITE-seq by mapping onto the CD4 MDE and testing candidate strategies by flow cytometry.

*Single-cell suspensions from spleen.* Single-cell suspensions were obtained by mechanical disruption through a 100 μm cell strainer, followed by red blood cell lysis using RBC lysis buffer (Thermo Fisher, Cat. #00-4333-57) for 3 min at room temperature.

*Single-cell suspensions from lung*: The right heart was perfused with 10 mL of cold PBS. Lungs were harvested, minced, and incubated in 25 mL of RPMI containing 1 mg/mL collagenase IV (Gibco Thermo Fisher Cat. #17104019) and 150 μg/mL DNase I (Sigma Aldrich SKU #10104159001) at 37 °C for 40 min with stirring (500 rpm). Digested tissues were passed through a 100 μm cell strainer, washed in PBS, and red blood cell were lysed using RBC lysis buffer for 3 min at room temperature.

*Single-cell suspensions from colon.* The colon was mechanically cleaned in cold PBS to remove fecal content, opened longitudinally, and cut into three pieces. Tissue was incubated in 25 mL of chemical dissociation medium (RPMI, 5% FBS, 1 mM DTT, and 20 mM EDTA) at 37 °C for 15 min on a magnetic stirrer (500 rpm) to remove epithelium and mucus. The remaining pieces were then minced and incubated in 25 mL of enzymatic digestion medium (RPMI, 1% FBS, 0.5 mg/mL Dispase II (Sigma-Aldrich, Cat. No. D4693-1G), and 1.5 mg/mL Collagenase II (Gibco, Thermo Fisher, Cat. No. 17101015)) for 40 min at 37 °C. The resulting suspension was homogenized by resuspension and filtered through a 40 μm strainer.

*Staining.* Up to 6 × 10^6^ cells were stained. Cells were first incubated with Zombie Aqua viability dye (BioLegend, Cat. #423101, 1:1000) for 20 min at 4 °C protected from light. After washing in FACS buffer (PBS, 1 mM EDTA, 2% FBS), cells were incubated with Fc Block (BD Biosciences, Cat. #553142) for 10 min, followed by surface antibody staining for 20 min at 4 °C in FACS buffer against CD62L (clone MEL-14, BUV737, BD Biosciences Cat. #612833, 1:400), CD44(clone IM7, BV785, Biolegend Cat No. 103059, 1:400), CXCR6 (clone SA051D1, APC, Biolegend Cat. #151106, 1:200), CD185/CXCR5 (clone L138D7, BV605, Biolegend Cat. #145513, 1:300), CD279/PD1 (clone 29F.1A12, BD Biosciences Cat. #568608, 1:400), FR4 (clone 12A5, PE, Biolegend Cat. #125007, 1:400), CD73 (clone TY/11.8, Alexa Fluor 700, Biolegend Cat. #127230, 1:200), IL-33R/ST2 (clone U29-93, BUV615, BD Biosciences Cat. #751293, 1:400), IL-23R (clone 12B2B64, BV421, Biolegend Cat. #150907, 1:100), Ly-6C (clone HK1.4, BUV563, BD Biosciences Cat. #755198, 1:400), KLRG1 (clone 2F1, BV711, Biolegend Cat. #138427, 1:300), TCRβ (clone H57-597, BV570, Biolegend Cat. #109231, 1:400), CD4 (clone GK1.4, PerCP, Biolegend Cat. #100432, 1:400), and CD8b (clone H35-17.2, BV750, Biolegend Cat. #423101, 1:400) (**Extended Data Table 5**). Brilliant Stain Buffer (Thermo Fisher, Cat. #00-440-942) was used in the staining solution when more than two BV-conjugated antibodies were included. For intracellular staining, cells were fixed with Foxp3 Fix/Perm buffer (Thermo Fisher, Cat. #00-5523-00) following the manufacturer’s protocol, for 30 min at 4 °C in the dark, washed in Perm/Wash buffer, and stained in 200 μL with antibodies against RORγt (clone B2D, PerCP-eFluor710, Thermo Fisher, Cat. #46-6981-82, 1:80), FoxP3 (clone MF-14, Alexa Fluor 488, BioLegend, Cat. #126406, 1:200), GATA3 (clone TWAJ, PE-eFluor610, Thermo Fisher, Cat. #61-9966-42, 1:80), and T-bet (1clone 4B10, PE-Fire810, BioLegend, Cat. #644839, 1:100) for 45 min at room temperature in the dark.

*Data acquisition and analysis.* on a Cytek Aurora (Cytek Biosciences). Data were analyzed using FlowJo v10 (TreeStar, BD LifeSciences).

### Plotting

Plots were generated in R using ggplot2^126^, S-Plus, Seurat^117^, or the ZemmourLib R package (https://github.com/dzemmour/ZemmourLib, v0.1.3). Heatmaps were generated using pheatmap (v1.0.13) or Morpheus (Broad Institute)

## Code availability

Code is available at the following repository:

https://github.com/immgen/immgenT_Project/tree/main/CD4_paper

## Data availability

All raw and processed sequencing data generated in this study are available through the Gene Expression Omnibus (GEO) under accession GSE297097.

## Studies mapped using T-RBI: GSE184259, GSE315686, GSE281547, GSE221337, GSE233703, GSE298371immgenT resource: public data browsers

Beyond the data and analyses reported in this and accompanying articles, immgenT’s goal as a public resource is served by a set of databrowsers which enable multi-faceted exploration of the data. They are accessible through the immgenT portal (https://www.immgen.org/ImmGenT/) and include:

**Rosetta2** (https://rosetta.immgen.org/) is an interactive single-cell viewer that connects RNA-seq and protein (CITE-seq) data. Cell populations can be visualized on parallel RNA and protein UMAPs, coloring reciprocally RNA- or protein-based clusters. The expression of individual genes, or of user defined gene signatures can be mapped on these representations. A cell gating interface modeled after flow cytometry serially generates 2D CITE-seq plots, on which cells can be “gated”, for projection onto UMAPs or computation of differential expression. Rosetta2 returns differentially expressed genes and proteins (as heatmaps or tables). Rosetta2 can be used to explore individual datasets or, or integrated cells grouped by lineage.

**TCR Browser** (https://rstats.immgen.org/tcrbrowser/) is a searchable interface for TCR sequences across the immgenT data. TCR repertoires can be browsed within immgenT datasets, searched for TCRs that use specific V, J or CDR3 elements, or for similarity to a user provided TCR.

**T-RBI portal** (Reference Based Integration) (https://rstats.immgen.org/immgenT_integration_analysis/) allows AI-driven mapping of external single-cel RNAseq datasets into the immgenT framework. Users upload their own data and metadata files (standard sc formats), which are then mapped and annotated within the immgenT framework. After an overnight compute, the tool returns annotation tables with integration statistics and annotation of every cell in the immgenT cluster framework, and x/y coordinates in the reference MDE space.

## Skyline browser

(https://rstats.immgen.org/Skyline/skyline.html?datagroup=immgenT%20Pseudobulk) represents, as a classic bar histogram, a pseudo-bulked overview of expression profiles of a given gene across all or selected immgenT cell clusters

---

**Extended Data Figure 1.**
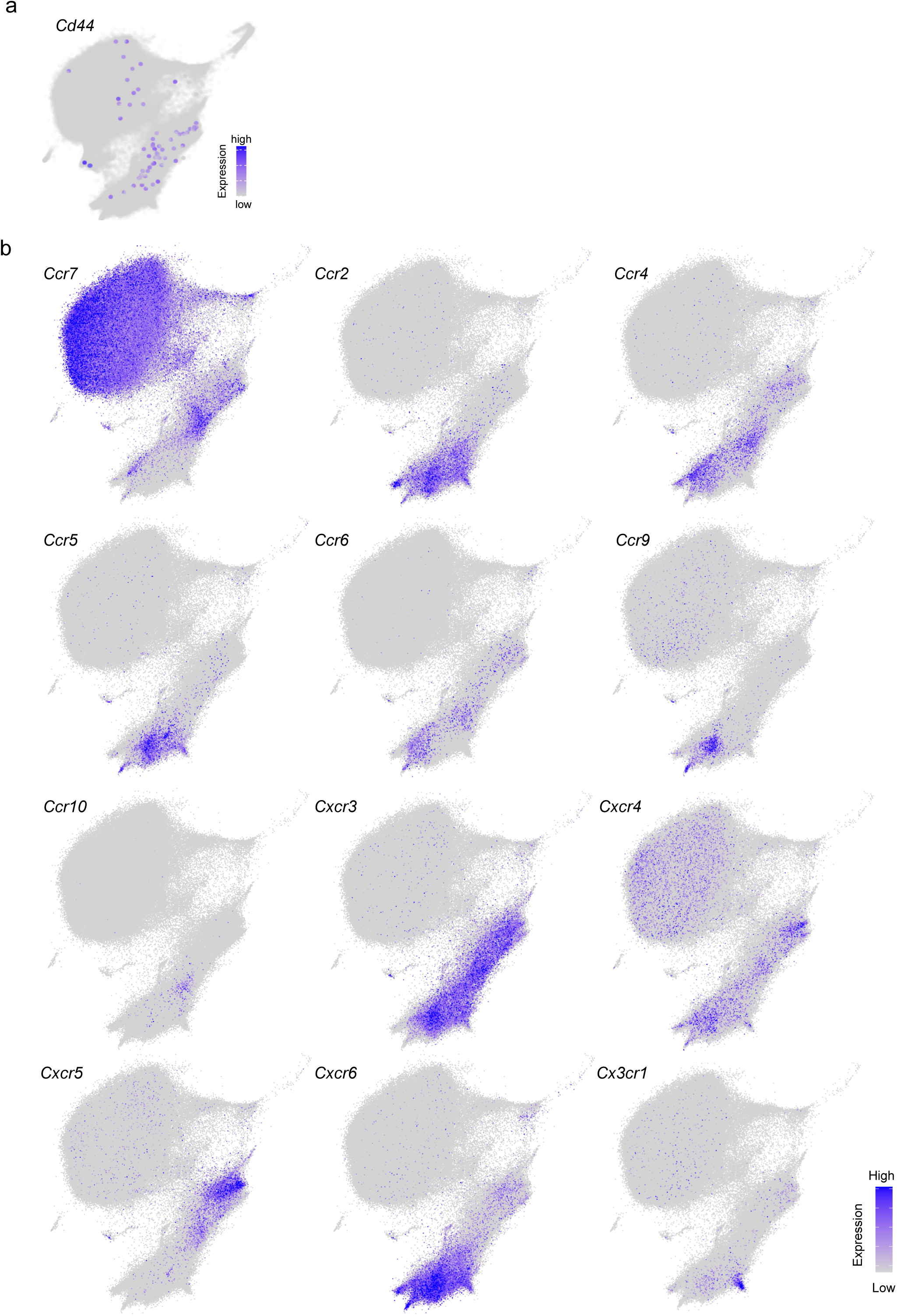
Chemokine receptor expression across the CD4 landscape. **(a)** Feature plot showing *Cd44* expression in antigen-specific memory CD4⁺ T cells following LCMV or *Listeria monocytogenes* infection in secondary lymphoid organs (SLOs). Background immgenT CD4⁺ T cells are shown in grey. **(b)** CD4 MDE showing expression of chemokine receptor genes.

**Extended Data Figure 2.**
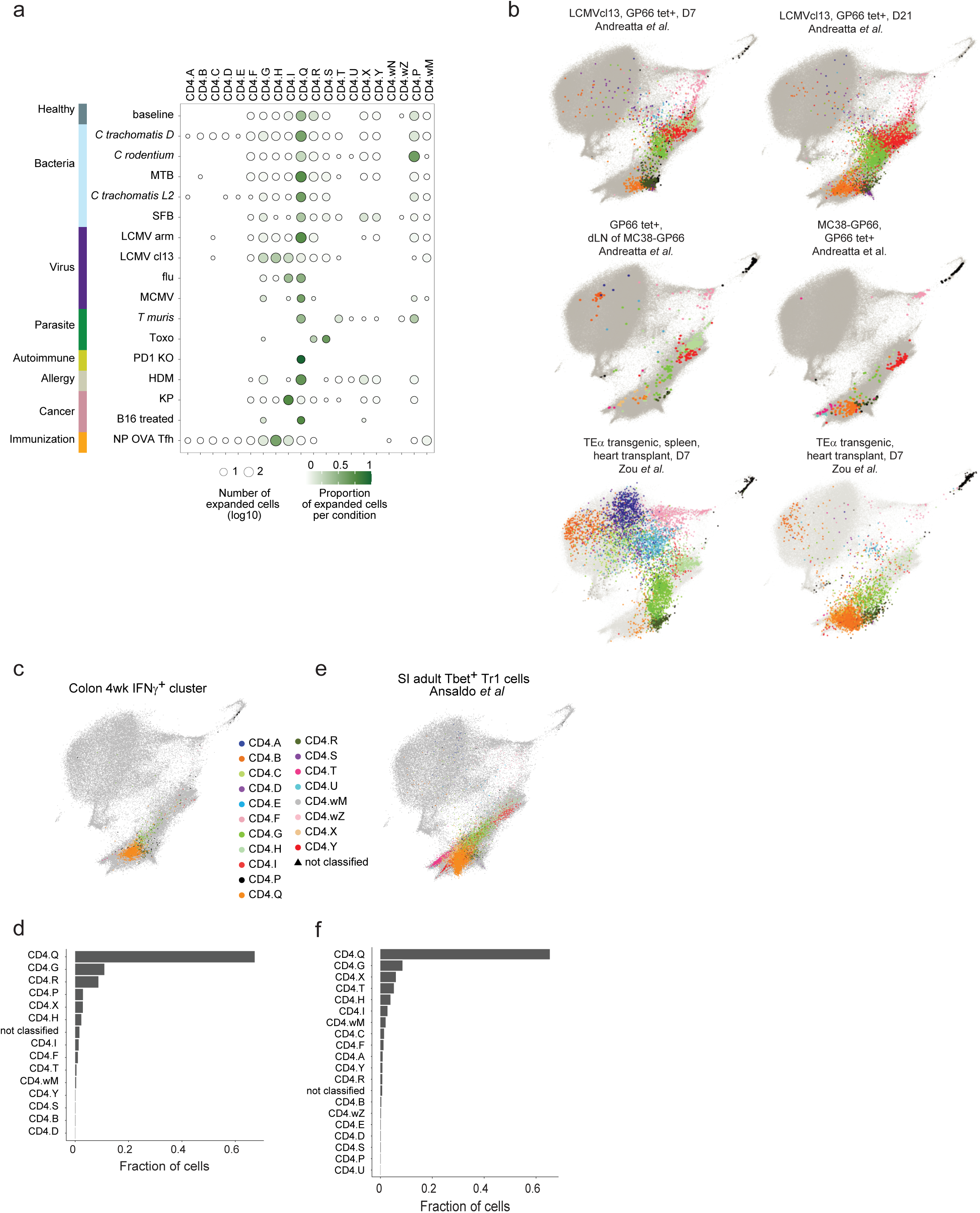
Cluster distribution of antigen-specific and clonally expanded cells across immune conditions. **(a)** Dot plot, similar to Fig. 1e, showing the distribution of clonally expanded cells (clonotypes containing ≥10 cells) across clusters. For each immune condition (row), dot size indicates the number of cells, and color indicates the mean proportion of cells in each cluster. Clonotypes were defined by unique paired αβ TCR sequences, including V-gene, J-gene and CDR3 nucleotide sequences. **(b)** CD4 MDE plots showing antigen-specific CD4⁺ T cells from published datasets projected onto the immgenT CD4 reference using T-RBI. Datasets included GP66 tetramer⁺ cells collected 7 and 21 days after LCMV clone 13 infection (GSE182320; Andreatta et al.^41^), GP66 tetramer⁺ cells from draining lymph nodes and MC38-GP tumours (GSE182320; Andreatta et al.^41^), and TEa transgenic cells from the spleen and heart after cardiac transplantation (GSE221337; Zou et al.^42^). Projected cells are colored according to their assigned immgenT CD4 cluster identity; background immgenT CD4 cells are shown in grey. **(c–f)** Examples illustrating the use of T-RBI to compare related cell populations generated by independent laboratories. **(c,d)** CD4 MDE **(c)** and cluster distribution **(d)** showing the projection of IFNγ⁺ CD4⁺ T cells isolated from the colon of 4-week-old mice in one of the authors’ laboratories. **(e,f)** CD4 MDE **(e)** and cluster distribution **(f)** showing the projection of T-bet⁺ CD4⁺ T cells isolated from the small intestine of adult mice and annotated as Tr1 cells by Ansaldo et al^43^. Projected cells are colored according to their assigned immgenT CD4 cluster identity; background immgenT CD4 cells are shown in grey.

**Extended Data Figure 3.**
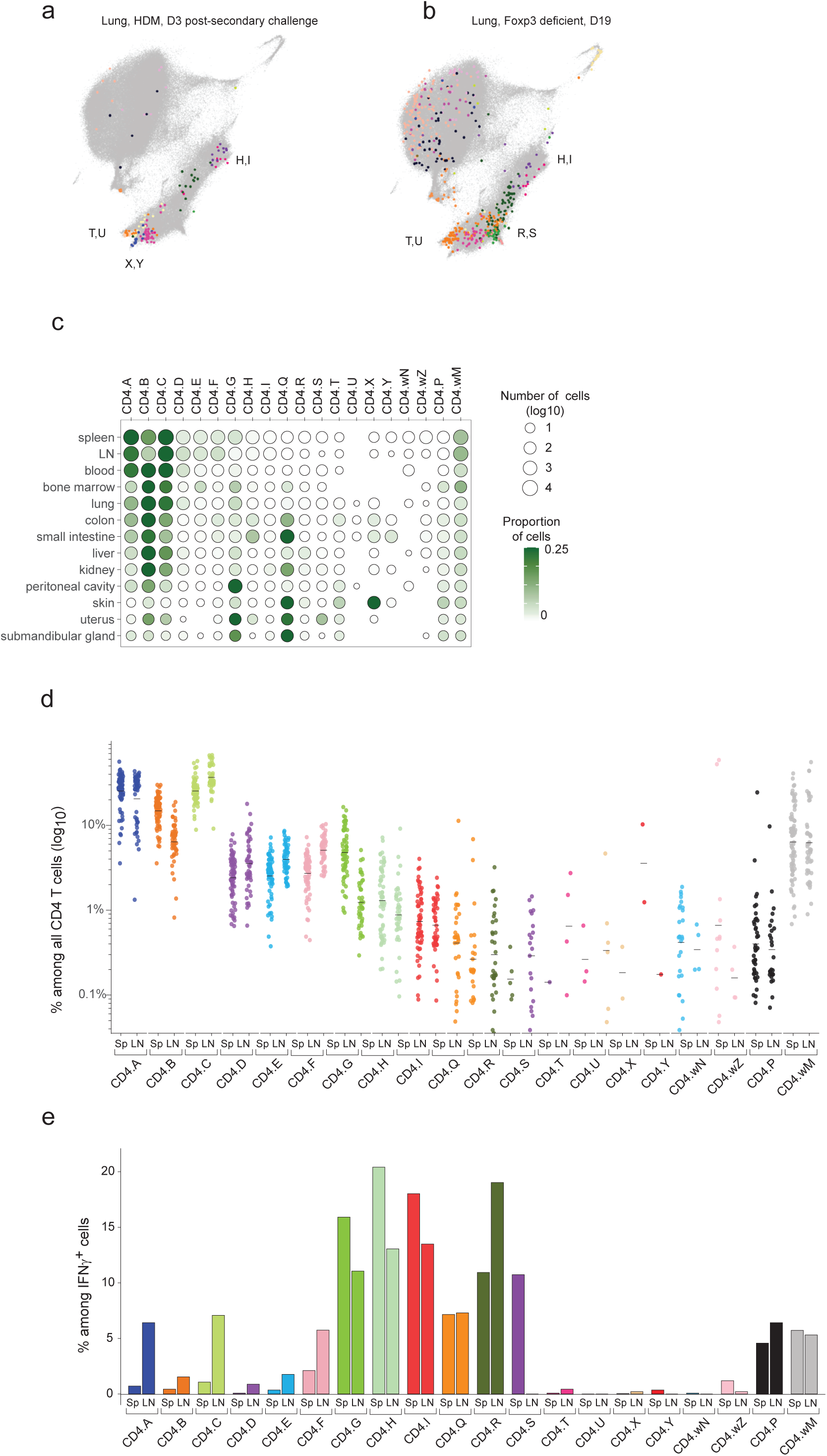
Polarized clusters co-occur within individual immune responses and exhibit tissue-specific biases. **(a,b)** CD4 MDE highlighting examples in which multiple Th polarized clusters co-occur within individual immune responses. **(a)** CD4 MDE showing house dust mite–specific CD4⁺ T cells (Der p1 tetramer⁺) in the lung at day 3 post-secondary challenge, a Th2 allergy model (IGT81). Cells are colored by cluster identity; background immgenT CD4+ T cells are shown in grey. **(b)** CD4 MDE showing CD4+ T cells in the lung of Foxp3-deficient mice at day 19 (IGT24), at the peak of systemic inflammation. Cells are colored by cluster identity; background immgenT CD4+ T cells are shown in grey. **(c)** Dot plot, similar to Fig. 1e, showing the proportions of CD4⁺ T cell clusters across tissues at baseline. **(d)** Scatterplot showing the proportions of CD4⁺ T cell clusters in the spleen and lymph nodes at baseline. Each point represents one sample; black horizontal bars indicate the mean. **(e)** Bar plot showing the cluster distribution of *Ifng*-expressing cells in the spleen and lymph nodes at baseline.

**Extended Data Figure 4.**
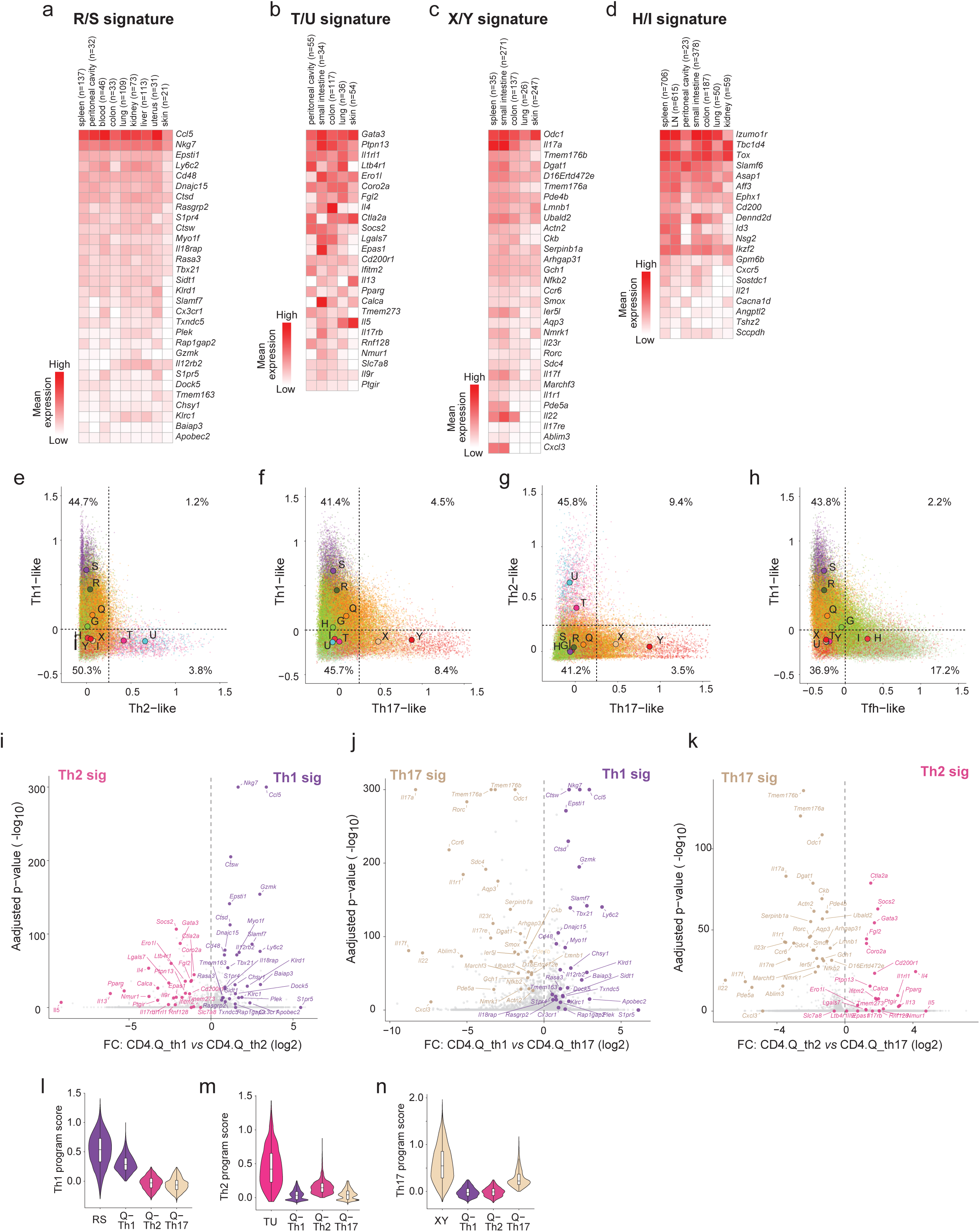
In vivo Th signature expression is conserved across tissues and varies in intensity and combination across clusters at the single-cell level. **(a–d)** Heatmaps showing expression of in vivo Th signature genes across tissues at baseline: **(a)** Th1 (R/S), **(b)** Th2 (T/U), **(c)** Th17 (X/Y), and **(d)** Tfh-like (H/I) signatures. Only tissues containing at least 20 cells are shown for each signature. Color intensity represents mean gene expression (log1p(CP10K)). **(e–h)** Scatterplots similar to Fig. 4f-i but showing pairwise expression of Th programs across all activated CD4 cells (CD4.G–Y). Larger labeled circles indicate mean program expression per cluster; percentages denote the fraction of cells in each quadrant. **(i–k)** Volcano plots showing differential gene expression between committed Q cells (Q-Th1, Q-Th2, and Q-Th17) and their respective polarized clusters. The plots show fold change versus adjusted *P* value. Th1, Th2, and Th17 signature genes, as defined in Fig. 4a–c, are highlighted. **(l–n)** Violin plots showing the distribution of Th1 (l), Th2 (m), and Th17 (n) signature expression in committed Q-Th1, Q-Th2, and Q-Th17 cells compared with the respective polarized clusters.

**Extended Data Figure 5.**
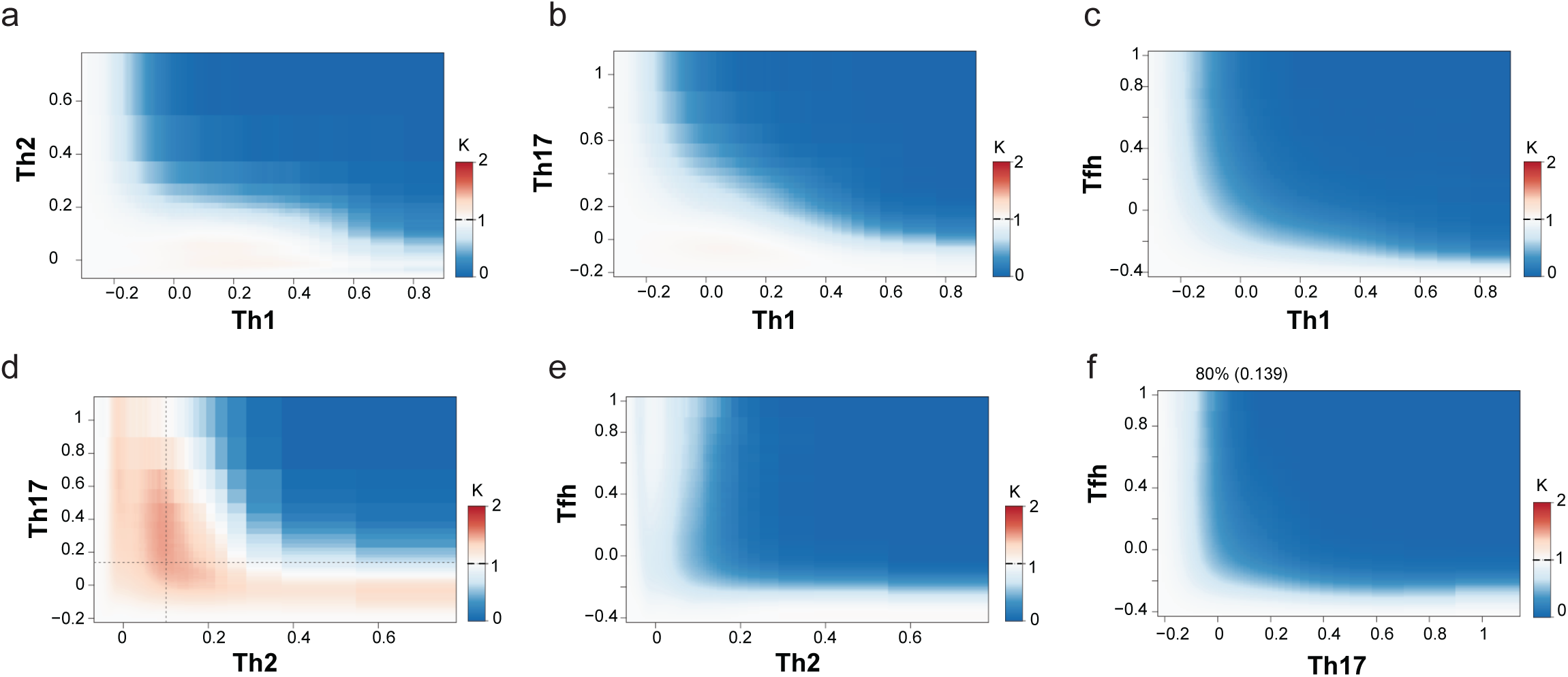
Polyfunctional cell frequencies reflect random assortment of Th programs at low expression levels. **(a–f)** Heatmaps showing the ratio of observed versus expected Th program co-expression. k(x,y) = Pr(THx>x, THy>y) / [Pr(THx>x)·Pr(THy>y)]. Values <1 (blue) indicate reduced co-expression than expected (mutual exclusion), observed at high Th expression. Values ≥1 (white–red) indicate no deviation from independence, consistent with random co-expression at lower Th expression levels (midland clusters).

**Extended Data Figure 6.**
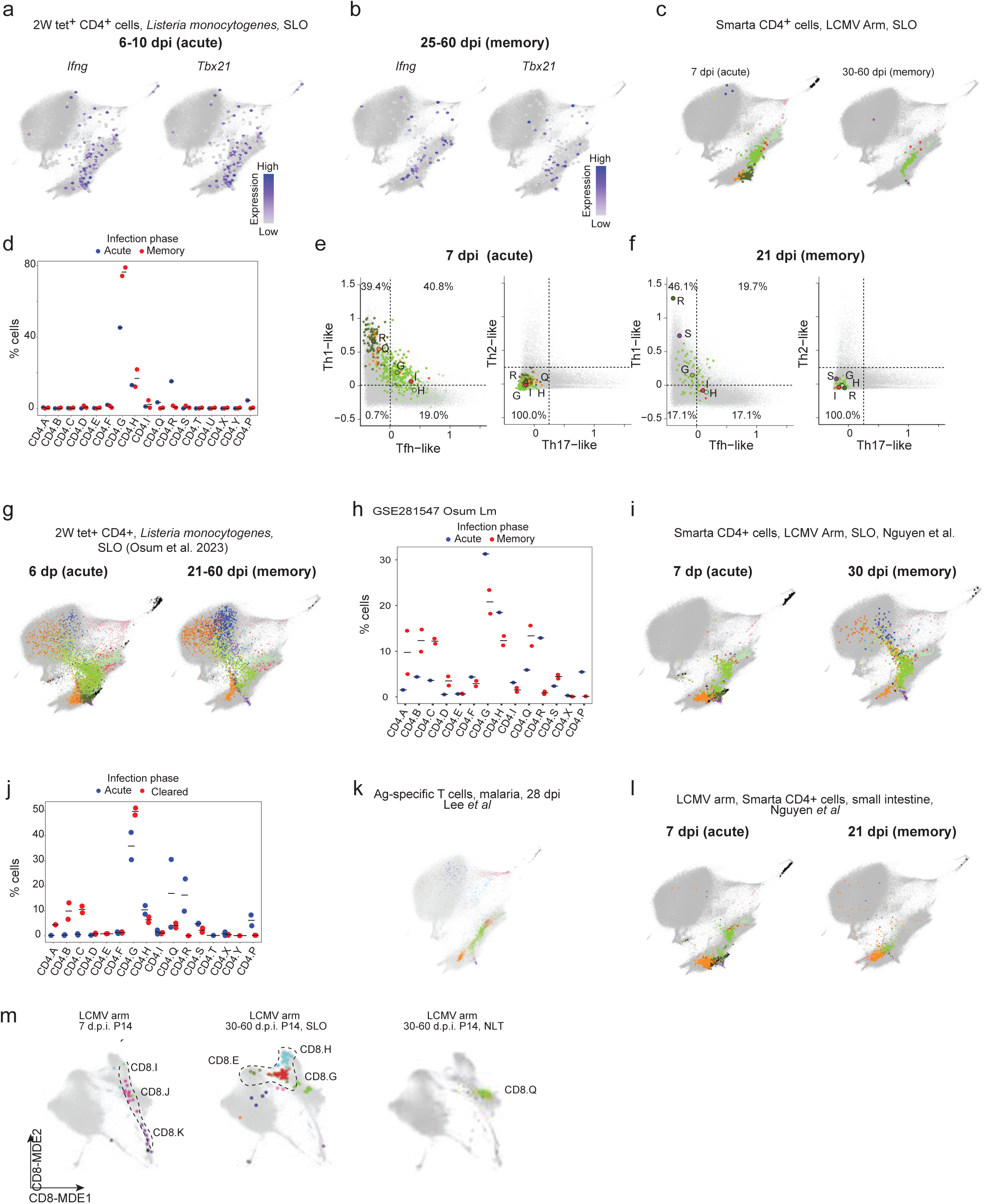
Memory CD4+ T cells. **(a,b)** CD4 MDE showing *Ifng* and *Tbx21* expression in *Listeria*-specific CD4⁺ T cells (2W tet⁺) in secondary lymphoid organs (SLO) at acute (6–10 dpi) (a) and memory (35–60 dpi) (b) (IGT43). 2W tet⁺ denotes cells recognizing *Listeria monocytogenes* engineered to express the 2W peptide. Background immgenT CD4+ T cells are shown in grey. **(c)** CD4 MDE showing the cluster distribution of LCMV-specific CD4⁺ T cells (SMARTA) in SLO at 7 dpi (acute, left) and 30–60 dpi (memory, right) (IGT36,38,40). Cells are colored by cluster identity; background immgenT CD4+ T cells are shown in grey. **(d)** Quantification from (c) showing cluster proportions at acute (blue) versus memory (red) phases in SLO. Each dot represents a sample; bars indicate the mean. **(e,f)** Scatterplots showing pairwise Th program expression in LCMV-specific CD4⁺ T cells at acute and memory phases. Larger labeled circles indicate the mean program expression per cluster (G-Y). **(g)** CD4 MDE showing projection of *Listeria*-specific CD4⁺ T cells (2W tet⁺) at acute (6 dpi) and memory (21-60 dpi) from Osum et al.^60^ onto the immgenT CD4 reference using T-RBI. Cells are colored by cluster identity; background immgenT CD4+ T cells are shown in grey (n = 4,500 cells per plot). **(h)** Quantification from (g) showing cluster proportions at acute (blue) versus memory (red) phases in SLO. Each dot represents a sample; bars indicate the mean. **(i)** CD4 MDE showing projection of LCMV-specific CD4⁺ T cells (SMARTA) at acute (7 dpi) and memory (21 dpi) in SLO from Nguyen et al.^69^ onto the immgenT CD4 reference using T-RBI. Cells are colored by cluster identity; background immgenT CD4+ T cells are shown in grey (n = 5,000 cells per plot). **(j)** Quantification from (i) showing cluster proportions at acute (blue) versus memory (red) phases in SLO. Each dot represents a sample; bars indicate the mean. **(k)** CD4 MDE showing the projection of malaria-specific CD4⁺ T cells 28 days after infection using T-RBI (GSE233703; Lee et al.^88^). Cells are colored according to their assigned immgenT CD4 cluster identity; background immgenT CD4 cells are shown in grey. **(l)** CD4 MDE showing projection of LCMV-specific CD4⁺ T cells (SMARTA) at acute (7 dpi) and memory (21 dpi) in the small intestine from Nguyen et al.^69^ onto the immgenT CD4 reference using T-RBI. Cells are colored by cluster identity; background immgenT CD4+ T cells are shown in grey (n = 1,800 cells per plot). **(m)** Unlike CD4⁺ T cells, CD8⁺ effector and memory cells occupy distinct clusters. immgenT CD8-specific MDE showing the distribution of LCMV-specific P14 CD8⁺ T cells at day 7 and at memory time points (30–60 dpi) following LCMV Armstrong infection. Memory P14 cells are shown separately for secondary lymphoid organs (SLOs) and non-lymphoid tissues (NLTs). Cells are colored according to their CD8 cluster identity; background cells from the immgenT CD8 reference are shown in grey.

**Extended Data Figure 7.**
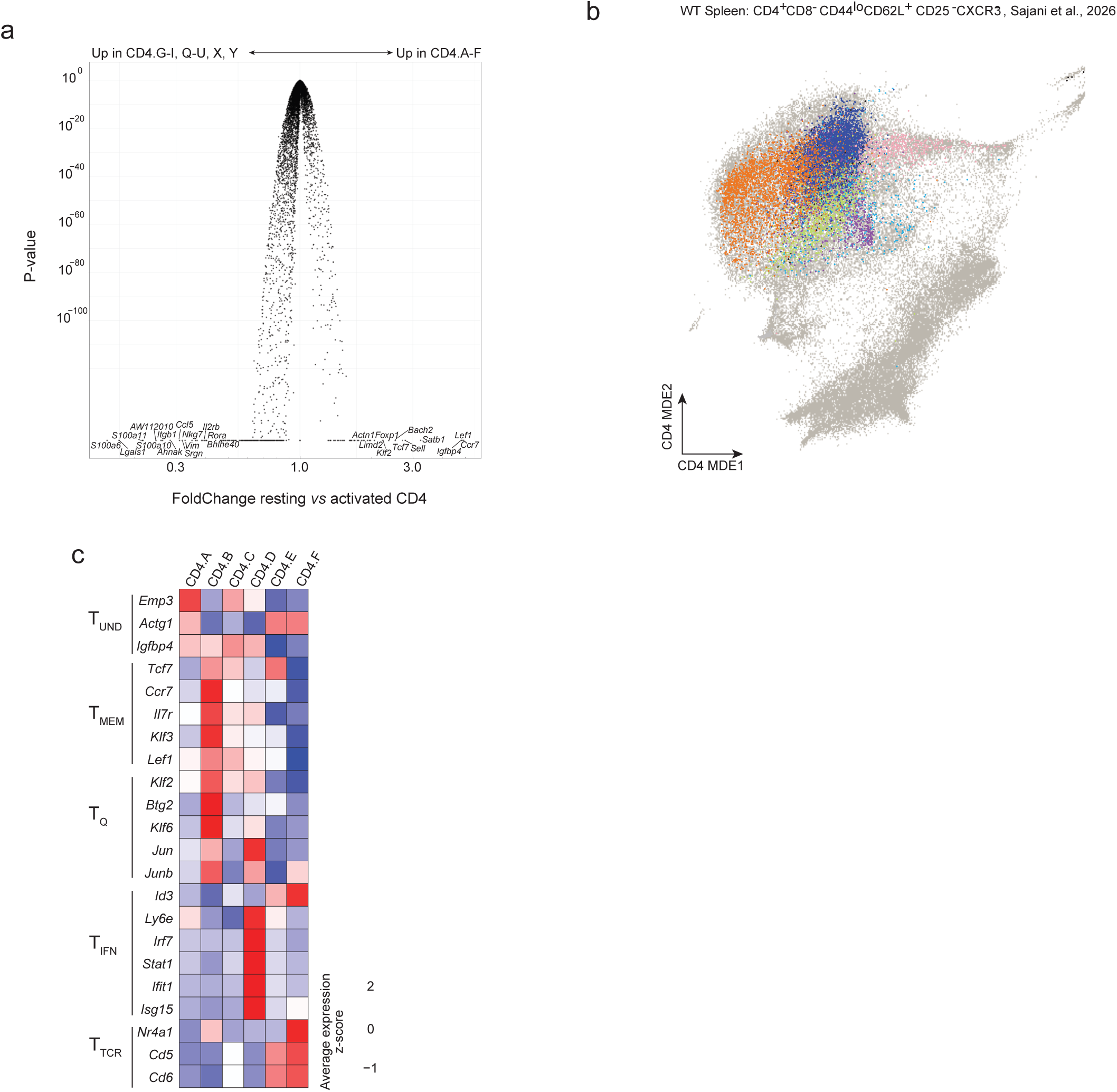
Resting CD4⁺ T cell clusters. **(a)** Volcano plot showing differential gene expression between resting (CD4.A–F) and activated (CD4.G–Y) clusters, plotted as fold change versus adjusted *P* value. **(b)** CD4 MDE showing projection of resting T cells from Sajani et al.^95^ onto the immgenT CD4 reference using T-RBI. Cells were sorted as CD4⁺CD8⁻CD44^lo^CD62L⁺CD25⁻CXCR3⁻ splenic cells at baseline. Cells are colored by cluster identity; background immgenT CD4+ T cells are shown in grey. **(c)** Heatmap showing expression of signature genes from five resting CD4⁺ T cell clusters identified by Sajani et al.^95^ (TUND, TMEM, TQ, TIFN, TTCR) across immgenT CD4.A–F clusters.

**Extended Data Figure 8.**
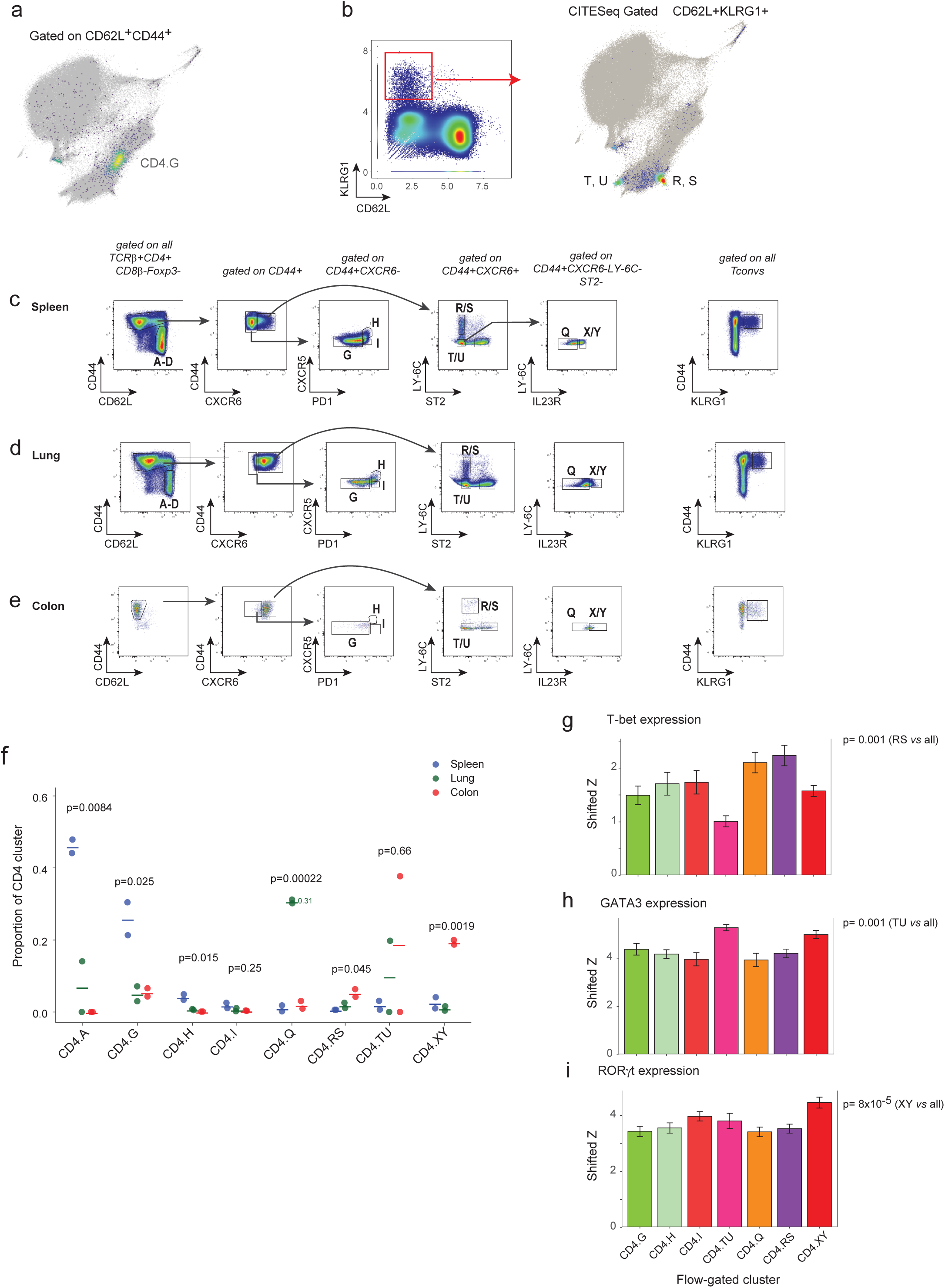
immgenT CD4 flow cytometry panel examples. **(a)** CD4 MDE highlighting cells bioinformatically gated (CITE-seq) as CD44⁺CD62L⁺, mapping to CD4.G. **(b)** CD4 MDE showing cells gated as CD62L⁻KLRG1⁺, mapping to Th1 (CD4.R,S) and Th2 (CD4.T,U) poles. Left: gating strategy showing CD62L versus KLRG1 (CITE-seq; log1p(CP10K)). **(c–e)** Representative flow cytometry plots showing the immgenT CD4 gating strategy in spleen, lung, and colon at baseline. **(f)** Proportion of indicated CD4 clusters within Tconv cells (TCRβ⁺CD4⁺CD8β⁻Foxp3⁻) across organs (n = 2); *P* value by ANOVA. **(g–i)** Relative expression of T-bet (g), GATA3 (h), and RORγt (i) across flow-gated CD4⁺ T cell clusters. Bars show means; error bars indicate SEM (n = 28 across 5 experiments). P value from two-sided t-test.

**Extended Data Table 1. Sample-level metadata for the immgenT dataset**

**Extended Data Table 2. Cluster signatures**

**Extended Data Table 3. Expanded TCR clonotypes across immune challenges**

**Extended Data Table 4. In vivo helper T cell signatures**

**Extended Data Table 5. CD4 flow cytometry panel**

**Extended Data Table 6. Correspondence between immgenT clusters and common annotations**

## References

1. Kisielow, P. et al. Ly antigens as markers for functionally distinct subpopulations of thymus-derived lymphocytes of the mouse. Nature 253, 219–220 (1975).

2. Cantor, H. & Boyse, E. A. Functional subclasses of T-lymphocytes bearing different Ly antigens. I. The generation of functionally distinct T-cell subclasses is a differentiative process independent of antigen. J. Exp. Med. 141, 1376–1389 (1975).

3. Swain, S. L. & Panfili, P. R. Helper cells activated by allogeneic H-2K or H-2D differences have a Ly phenotype distinct from those responsive to I differences. J. Immunol. 122, 383–391 (1979).

4. Laidlaw, B. J., Craft, J. E. & Kaech, S. M. The multifaceted role of CD4(+) T cells in CD8(+) T cell memory. Nat. Rev. Immunol. 16, 102–111 (2016).

5. Crotty, S. Follicular helper CD4 T cells (TFH). Annu. Rev. Immunol. 29, 621–663 (2011).

6. Swain, S. L., McKinstry, K. K. & Strutt, T. M. Expanding roles for CD4^+^ T cells in immunity to viruses. Nat. Rev. Immunol. 12, 136–148 (2012).

7. Ruterbusch, M., Pruner, K. B., Shehata, L. & Pepper, M. In vivo CD4+ T cell differentiation and function: Revisiting the Th1/Th2 paradigm. Annu. Rev. Immunol. 38, 705–725 (2020).

8. Tubo, N. J. & Jenkins, M. K. CD4+ T Cells: guardians of the phagosome. Clin. Microbiol. Rev. 27, 200–213 (2014).

9. Barré-Sinoussi, F. et al. Isolation of a T-lymphotropic retrovirus from a patient at risk for acquired immune deficiency syndrome (AIDS). Science 220, 868–871 (1983).

10. Kabat, A. M. et al. Resident TH2 cells orchestrate adipose tissue remodeling at a site adjacent to infection. Sci. Immunol. 7, eadd3263 (2022).

11. Boothby, I. C. et al. Early-life inflammation primes a T helper 2 cell-fibroblast niche in skin. Nature 599, 667–672 (2021).

12. Jaquish, A. et al. Mammary intraepithelial lymphocytes and intestinal inputs shape T cell dynamics in lactogenesis. Nat. Immunol. 26, 1411–1422 (2025).

13. Mosmann, T. R., Cherwinski, H., Bond, M. W., Giedlin, M. A. & Coffman, R. L. Two types of murine helper T cell clone. I. Definition according to profiles of lymphokine activities and secreted proteins. J. Immunol. 136, 2348–2357 (1986).

14. Schnell, A., Littman, D. R. & Kuchroo, V. K. TH17 cell heterogeneity and its role in tissue inflammation. Nat. Immunol. 24, 19–29 (2023).

15. Liew, F. Y. T(H)1 and T(H)2 cells: a historical perspective. Nat. Rev. Immunol. 2, 55–60 (2002).

16. Korn, T., Bettelli, E., Oukka, M. & Kuchroo, V. K. IL-17 and Th17 cells. Annu. Rev. Immunol. 27, 485–517 (2009).

17. Zhu, J., Yamane, H. & Paul, W. E. Differentiation of effector CD4 T cell populations (*). Annu. Rev. Immunol. 28, 445–489 (2010).

18. Butcher, M. J. & Zhu, J. Recent advances in understanding the Th1/Th2 effector choice. Fac. Rev. 10, 30 (2021).

19. Vivier, E. et al. Innate lymphoid cells: 10 years on. Cell 174, 1054–1066 (2018).

20. Jameson, S. C. & Masopust, D. Understanding Subset Diversity in T Cell Memory. Immunity 48, 214–226 (2018).

21. Kelso, A. Th1 and Th2 subsets: paradigms lost? Immunol. Today 16, 374–379 (1995).

22. O’Shea, J. J. & Paul, W. E. Mechanisms underlying lineage commitment and plasticity of helper CD4+ T cells. Science 327, 1098–1102 (2010).

23. Bluestone, J. A., Mackay, C. R., O’Shea, J. J. & Stockinger, B. The functional plasticity of T cell subsets. Nat. Rev. Immunol. 9, 811–816 (2009).

24. Rogozynski, N. P. & Dixon, B. The Th1/Th2 paradigm: A misrepresentation of helper T cell plasticity. Immunol. Lett. 268, 106870 (2024).

25. Oestreich, K. J. & Weinmann, A. S. Master regulators or lineage-specifying? Changing views on CD4+ T cell transcription factors. Nat. Rev. Immunol. **12**, 799–804 (2012).

26. Gaublomme, J. T. et al. Single-Cell Genomics Unveils Critical Regulators of Th17 Cell Pathogenicity. Cell 163, 1400–1412 (2015).

27. Eizenberg-Magar, I. et al. Diverse continuum of CD4+ T-cell states is determined by hierarchical additive integration of cytokine signals. Proc. Natl. Acad. Sci. U. S. A. 114, E6447–E6456 (2017).

28. Kiner, E. et al. Gut CD4+ T cell phenotypes are a continuum molded by microbes, not by TH archetypes. Nat. Immunol. 22, 216–228 (2021).

29. Magill, I., et al. ImmgenT: A comprehensive reference of convergent T-cell states in the mouse. *bioRxiv* 2026.01. 30.702892 (2026) doi:10.64898/2026.01.30.702892.

30. Gayoso, A. et al. Joint probabilistic modeling of single-cell multi-omic data with totalVI. Nat. Methods 18, 272–282 (2021).

31. Sallusto, F., Lenig, D., Mackay, C. R. & Lanzavecchia, A. Flexible programs of chemokine receptor expression on human polarized T helper 1 and 2 lymphocytes. J. Exp. Med. 187, 875–883 (1998).

32. Singh, S. P., Zhang, H. H., Foley, J. F., Hedrick, M. N. & Farber, J. M. Human T cells that are able to produce IL-17 express the chemokine receptor CCR6. J. Immunol. 180, 214–221 (2008).

33. Croze, M., et al. The αβTCR repertoire at scale in the immgenT dataset. bioRxiv (2026) doi:10.64898/2026.01.30.702900.

34. Milighetti, M. et al. Large clones of pre-existing T cells drive early immunity against SARS-COV-2 and LCMV infection. iScience 26, 106937 (2023).

35. Minervina, A. A. et al. Primary and secondary anti-viral response captured by the dynamics and phenotype of individual T cell clones. Elife 9, e53704 (2020).

36. Oliveira, G. et al. Phenotype, specificity and avidity of antitumour CD8+ T cells in melanoma. Nature 596, 119–125 (2021).

37. Zhou, Y. D. et al. T cell fate is dictated by different antigen-presenting cells in response to dietary versus gut epithelial self-antigen. Immunity 0, (2026).

38. Chi, X. et al. TCR origin and specificity to environmental or self-antigens on distinct antigen-presenting cells determine peripheral Treg cell differentiation. Immunity 0, (2026).

39. Lopez, R., Regier, J., Cole, M. B., Jordan, M. I. & Yosef, N. Deep generative modeling for single-cell transcriptomics. Nat. Methods 15, 1053–1058 (2018).

40. Xu, C. et al. Probabilistic harmonization and annotation of single-cell transcriptomics data with deep generative models. Mol. Syst. Biol. 17, e9620 (2021).

41. Andreatta, M. et al. A CD4+ T cell reference map delineates subtype-specific adaptation during acute and chronic viral infections. Elife 11, (2022).

42. Zou, D. et al. CD4+ T cell immunity is dependent on an intrinsic stem-like program. Nat. Immunol. 25, 66–76 (2024).

43. Ansaldo, E. et al. T-bet-expressing Tr1 cells driven by dietary signals dominate the small intestinal immune landscape. Proc. Natl. Acad. Sci. U. S. A. 123, e2520747122 (2026).

44. Kawabe, T. et al. Memory-phenotype CD4+ T cells spontaneously generated under steady-state conditions exert innate TH1-like effector function. Sci. Immunol. 2, eaam9304 (2017).

45. Chi, X. et al. RORγt expression in mature TH17 cells safeguards their lineage specification by inhibiting conversion to TH2 cells. Sci. Adv. 8, eabn7774 (2022).

46. Drujont, L. et al. RORγt+ cells selectively express redundant cation channels linked to the Golgi apparatus. Sci. Rep. 6, 23682 (2016).

47. Ciofani, M. et al. A validated regulatory network for Th17 cell specification. Cell 151, 289–303 (2012).

48. Wynn, T. A. Type 2 cytokines: mechanisms and therapeutic strategies. Nat. Rev. Immunol. 15, 271–282 (2015).

49. Yu, D. et al. The transcriptional repressor Bcl-6 directs T follicular helper cell lineage commitment. Immunity 31, 457–468 (2009).

50. Nurieva, R. I. et al. Generation of T follicular helper cells is mediated by interleukin-21 but independent of T helper 1, 2, or 17 cell lineages. Immunity 29, 138–149 (2008).

51. Hsieh, C. S. et al. Development of TH1 CD4+ T cells through IL-12 produced by Listeria-induced macrophages. Science 260, 547–549 (1993).

52. Jarick, K. J. et al. Non-redundant functions of group 2 innate lymphoid cells. Nature 611, 794–800 (2022).

53. Cannons, J. L., Tangye, S. G. & Schwartzberg, P. L. SLAM family receptors and SAP adaptors in immunity. Annu. Rev. Immunol. 29, 665–705 (2011).

54. Xiao, S. et al. Small-molecule RORγt antagonists inhibit T helper 17 cell transcriptional network by divergent mechanisms. Immunity 40, 477–489 (2014).

55. Nguyen, C. et al. Bhlhe40 promotes CD4+ T helper 1 cell and suppresses T follicular helper cell differentiation during viral infection. J. Immunol. 212, 1829–1842 (2024).

56. Galletti, G. et al. The CD8 immgenT framework as a universal reference of mouse CD8αβ T cell differentiation states. *bioRxivorg* 2026.02.02.703365 (2026) doi:10.64898/2026.02.02.703365.

57. Cook, M. E., Jarjour, N. N., Lin, C.-C. & Edelson, B. T. Transcription factor Bhlhe40 in immunity and autoimmunity. Trends Immunol. 41, 1023–1036 (2020).

58. Carlson, C. M. et al. Kruppel-like factor 2 regulates thymocyte and T-cell migration. Nature 442, 299–302 (2006).

59. Scott, M. C. et al. Deep profiling deconstructs features associated with memory CD8+ T cell tissue residence. Immunity 58, 162–181.e10 (2025).

60. Osum, K. C. et al. A minority of Th1 and Tfh effector cells express survival genes shared by memory cell progeny that require IL-7 or TCR signaling to persist. Cell Rep. 44, 115111 (2025).

61. Groux, H. et al. A CD4+ T-cell subset inhibits antigen-specific T-cell responses and prevents colitis. Nature 389, 737–742 (1997).

62. Trifari, S., Kaplan, C. D., Tran, E. H., Crellin, N. K. & Spits, H. Identification of a human helper T cell population that has abundant production of interleukin 22 and is distinct from T(H)-17, T(H)1 and T(H)2 cells. Nat. Immunol. 10, 864–871 (2009).

63. Duhen, T., Geiger, R., Jarrossay, D., Lanzavecchia, A. & Sallusto, F. Production of interleukin 22 but not interleukin 17 by a subset of human skin-homing memory T cells. Nat. Immunol. 10, 857–863 (2009).

64. Ahlfors, H. et al. IL-22 fate reporter reveals origin and control of IL-22 production in homeostasis and infection. J. Immunol. 193, 4602–4613 (2014).

65. Barnes, J. L. et al. T-helper 22 cells develop as a distinct lineage from Th17 cells during bacterial infection and phenotypic stability is regulated by T-bet. Mucosal Immunol. 14, 1077–1087 (2021).

66. McDonald, P. W. et al. IL-7 signalling represses Bcl-6 and the TFH gene program. Nat. Commun. 7, 10285 (2016).

67. Xia, Y. et al. BCL6-dependent TCF-1+ progenitor cells maintain effector and helper CD4+ T cell responses to persistent antigen. Immunity 55, 1200–1215.e6 (2022).

68. Wu, T., et al. The TCF1-Bcl6 axis counteracts type I interferon to repress exhaustion and maintain T cell stemness. Sci. Immunol. 1, eaai8593 (2016).

69. Nguyen, Q. P. et al. Transcriptional programming of CD4+ TRM differentiation in viral infection balances effector- and memory-associated gene expression. Sci. Immunol. 8, eabq7486 (2023).

70. Greczmiel, U. et al. Sustained T follicular helper cell response is essential for control of chronic viral infection. Sci. Immunol. 2, eaam8686 (2017).

71. Vinuesa, C. G., Linterman, M. A., Yu, D. & MacLennan, I. C. M. Follicular helper T cells. Annu. Rev. Immunol. 34, 335–368 (2016).

72. Clement, R. L. et al. Follicular regulatory T cells control humoral and allergic immunity by restraining early B cell responses. Nat. Immunol. 20, 1360–1371 (2019).

73. Shaw, L. A. et al. Id2 reinforces TH1 differentiation and inhibits E2A to repress TFH differentiation. Nat. Immunol. 17, 834–843 (2016).

74. Dangi, A., Husain, I., Jordan, C. Z., Yu, S. & Luo, X. Conversion of CD73hiFR4hi anergic T cells to IFN-γ-producing effector cells disrupts established immune tolerance. J. Clin. Invest. 133, e163872 (2023).

75. Kalekar, L. A. et al. CD4(+) T cell anergy prevents autoimmunity and generates regulatory T cell precursors. Nat. Immunol. 17, 304–314 (2016).

76. Iyer, S. S. et al. Identification of novel markers for mouse CD4(+) T follicular helper cells. Eur. J. Immunol. 43, 3219–3232 (2013).

77. Victora, G. D. & Nussenzweig, M. C. Germinal centers. Annu. Rev. Immunol. 40, 413–442 (2022).

78. Crotty, S. T follicular helper cell biology: A decade of discovery and diseases. Immunity 50, 1132–1148 (2019).

79. Marina-Zárate, E. et al. Highly functional and prolonged germinal center T follicular helper cell responses are associated with enhanced neutralizing antibody development. Immunity 58, 3094–3112.e7 (2025).

80. Sun, Q. et al. BCL6 promotes a stem-like CD8+ T cell program in cancer via antagonizing BLIMP1. Sci. Immunol. 8, eadh1306 (2023).

81. Osum, K. C. & Jenkins, M. K. Toward a general model of CD4+ T cell subset specification and memory cell formation. Immunity 56, 475–484 (2023).

82. Künzli, M. & Masopust, D. CD4+ T cell memory. Nat. Immunol. 24, 903–914 (2023).

83. Marshall, H. D. et al. Differential expression of Ly6C and T-bet distinguish effector and memory Th1 CD4(+) cell properties during viral infection. Immunity 35, 633–646 (2011).

84. Ahmed, R. & Gray, D. Immunological memory and protective immunity: understanding their relation. Science 272, 54–60 (1996).

85. Hale, J. S. et al. Distinct memory CD4+ T cells with commitment to T follicular helper- and T helper 1-cell lineages are generated after acute viral infection. Immunity 38, 805–817 (2013).

86. Khatun, A. et al. Single-cell lineage mapping of a diverse virus-specific naive CD4 T cell repertoire. J. Exp. Med. 218, (2021).

87. Shaw, L. A. et al. Id3 expression identifies CD4+ memory Th1 cells. Proc. Natl. Acad. Sci. U. S. A. 119, e2204254119 (2022).

88. Lee, H. J. et al. CD4+ T cells display a spectrum of recall dynamics during re-infection with malaria parasites. Nat. Commun. 15, 5497 (2024).

89. Osborne, L. C. et al. Impaired CD8 T cell memory and CD4 T cell primary responses in IL-7R alpha mutant mice. J. Exp. Med. 204, 619–631 (2007).

90. Wheeler, B. D., et al. The lncRNA Malat1 inhibits miR-15/16 to enhance cytotoxic T cell activation and memory cell formation. >bioRxivorg (2023) doi:10.1101/2023.04.14.536843.

91. Collins, N. et al. Skin CD4(+) memory T cells exhibit combined cluster-mediated retention and equilibration with the circulation. Nat. Commun. 7, 11514 (2016).

92. ElTanbouly, M. A. et al. VISTA is a checkpoint regulator for naïve T cell quiescence and peripheral tolerance. Science 367, eaay0524 (2020).

93. Even, Z. et al. The amalgam of naive CD4+ T cell transcriptional states is reconfigured by helminth infection to dampen the amplitude of the immune response. Immunity 57, 1893–1907.e6 (2024).

94. Deep, D. et al. Precursor central memory versus effector cell fate and naïve CD4+ T cell heterogeneity. J. Exp. Med. 221, (2024).

95. Sajani, A. et al. Heterogeneity and plasticity of the naive CD4+ T cell compartment. Cell Rep. 45, 116980 (2026).

96. Arazi, A. et al. The immune cell landscape in kidneys of patients with lupus nephritis. Nat. Immunol. 20, 902–914 (2019).

97. Tibbitt, C. A. et al. Single-Cell RNA Sequencing of the T Helper Cell Response to House Dust Mites Defines a Distinct Gene Expression Signature in Airway Th2 Cells. Immunity 10.1016/j.immuni.2019.05.014 (2019) doi:10.1016/j.immuni.2019.05.014.

98. Szabo, S. J. et al. A novel transcription factor, T-bet, directs Th1 lineage commitment. Cell 100, 655–669 (2000).

99. Zheng, W. & Flavell, R. A. The transcription factor GATA-3 is necessary and sufficient for Th2 cytokine gene expression in CD4 T cells. Cell 89, 587–596 (1997).

100. Harrington, L. E. et al. Interleukin 17-producing CD4+ effector T cells develop via a lineage distinct from the T helper type 1 and 2 lineages. Nat. Immunol. 6, 1123–1132 (2005).

101. Cano-Gamez, E. et al. Single-cell transcriptomics identifies an effectorness gradient shaping the response of CD4+ T cells to cytokines. Nat. Commun. 11, 1801 (2020).

102. Tubo, N. J. et al. Single naive CD4+ T cells from a diverse repertoire produce different effector cell types during infection. Cell 153, 785–796 (2013).

103. DiToro, D. et al. Differential IL-2 expression defines developmental fates of follicular versus nonfollicular helper T cells. Science 361, eaao2933 (2018).

104. Canesso, M. C. C. et al. Identification of antigen-presenting cell–T cell interactions driving immune responses to food. Science 10.1126/science.ado5088 (2024) doi:10.1126/science.ado5088.

105. Krishnamoorthy, V. et al. The IRF4 gene regulatory module functions as a read-write integrator to dynamically coordinate T helper cell fate. Immunity 47, 481–497.e7 (2017).

106. Krueger, P. D. et al. Two sequential activation modules control the differentiation of protective T helper-1 (Th1) cells. Immunity 54, 687–701.e4 (2021).

107. Lee, J.-Y. et al. The transcription factor KLF2 restrains CD4^+^ T follicular helper cell differentiation. Immunity 42, 252–264 (2015).

108. Pepper, M. et al. Different routes of bacterial infection induce long-lived TH1 memory cells and short-lived TH17 cells. Nat. Immunol. 11, 83–89 (2010).

109. Son, Y. M. et al. Tissue-resident CD4+ T helper cells assist the development of protective respiratory B and CD8+ T cell memory responses. Sci. Immunol. 6, eabb6852 (2021).

110. Swarnalekha, N. et al. T resident helper cells promote humoral responses in the lung. Sci. Immunol. 6, eabb6808 (2021).

111. Lee, V. et al. The endogenous repertoire harbors self-reactive CD4+ T cell clones that adopt a follicular helper T cell-like phenotype at steady state. Nat. Immunol. 24, 487–500 (2023).

112. Künzli, M. et al. Long-lived T follicular helper cells retain plasticity and help sustain humoral immunity. Sci. Immunol. 5, eaay5552 (2020).

113. Sallin, M. A. et al. Th1 differentiation drives the accumulation of intravascular, non-protective CD4 T cells during tuberculosis. Cell Rep. 18, 3091–3104 (2017).

114. Goldberg, M. F. et al. Salmonella persist in activated macrophages in T cell-sparse granulomas but are contained by surrounding CXCR3 ligand-positioned Th1 cells. Immunity 49, 1090–1102.e7 (2018).

115. Soon, M. S. F. et al. Transcriptome dynamics of CD4+ T cells during malaria maps gradual transit from effector to memory. Nat. Immunol. 21, 1597–1610 (2020).

116. Stoeckius, M. et al. Cell Hashing with barcoded antibodies enables multiplexing and doublet detection for single cell genomics. Genome Biol. 19, 224 (2018).

117. Satija, R. Tools for Single Cell Genomics [R package Seurat version 5.4.0]. Comprehensive R Archive Network (CRAN) https://CRAN.R-project.org/package=Seurat (2025).

118. Hao, Y. et al. Dictionary learning for integrative, multimodal and scalable single-cell analysis. Nat. Biotechnol. 10.1038/s41587-023-01767-y (2023) doi:10.1038/s41587-023-01767-y.

119. Luecken, M. D. et al. Benchmarking atlas-level data integration in single-cell genomics. Nat. Methods 19, 41–50 (2022).

120. Agrawal, A., Ali, A. & Boyd, S. Minimum-distortion embedding. Found. Trends® Mach. Learn. 14, 211–378 (2021).

121. Ritchie, M. E. et al. limma powers differential expression analyses for RNA-sequencing and microarray studies. Nucleic Acids Res. 43, e47 (2015).

122. Soneson, C. & Robinson, M. D. Bias, robustness and scalability in single-cell differential expression analysis. Nat. Methods 10.1038/nmeth.4612 (2018) doi:10.1038/nmeth.4612.

123. Liu, Y. et al. Dissecting tumor transcriptional heterogeneity from single-cell RNA-seq data by generalized binary covariance decomposition. Nat. Genet. 57, 263–273 (2025).

124. Empirical Bayes Matrix Factorization [R package flashier version 1.0.7]. Comprehensive R Archive Network (CRAN) https://CRAN.R-project.org/package=flashier (2023).

125. Delaney, C. et al. Combinatorial prediction of marker panels from single-cell transcriptomic data. Mol. Syst. Biol. 15, e9005 (2019).

126. Wickham, H. Ggplot2: Elegant Graphics for Data Analysis. (Springer New York, 2009).

